# Interplay between R2R3 MYB-type activators and repressors regulates proanthocyanidin biosynthesis in banana (*Musa acuminata*)

**DOI:** 10.1101/2022.04.27.489641

**Authors:** Ruchika Rajput, Jogindra Naik, Ralf Stracke, Ashutosh Pandey

## Abstract

- Proanthocyanidins are oligomeric flavonoid pigments that promote plant disease resistance and benefit human health. However, the transcriptional regulatory network that fine-tunes proanthocyanidin biosynthesis in banana (*Musa acuminata*) fruit remains poorly understood.
- We characterized two proanthocyanidin-specific R2R3 MYB activators (MaMYBPA1-MaMYBPA2) and four repressors (MaMYBPR1–MaMYBPR4) to elucidate the mechanisms underlying the transcriptional regulation of proanthocyanidin biosynthesis in banana.
- Heterologous expression of *MaMYBPA1* and *MaMYBPA2* partially complemented the *Arabidopsis thaliana* proanthocyanidin-deficient *transparent testa2* mutant. MaMYBPA1 and MaMYBPA2 interacted physically with MaMYCs and bound the promoters of the genes encoding anthocyanin synthase, leucoanthocyanidin reductase, and anthocyanidin reductase *in vitro* and form functional MBW complexes with MaTTG1 in *A. thaliana* protoplasts to transactivated these promoters *in vivo*.
- Overexpression of *MaMYBPA*s alone or with *MaMYC* in banana fruits induced proanthocyanidin accumulation and the transcription of proanthocyanidin biosynthesis– related genes. MaMYBPR repressors are also shown to interact with MaMYCs, forming repressing MBW complexes, and diminished proanthocyanidin accumulation. Interestingly the overexpression of MaMYBPA induces the expression of MaMYBPR, indicating an agile regulation of proanthocyanidin biosynthesis via the formation of competitive MBW complex. Taken together, our results reveal regulatory modules of R2R3 MYB- that fine-tune proanthocyanidin biosynthesis and offer possible targets for genetic manipulation in banana.

## Introduction

Proanthocyanidins are oligomeric flavonoids that accumulate in seeds, fruits, and vegetative tissues (Cipollini and Stiles, 1993; Barbehenn and Constabel, 2011). They produce seed coat pigmentation, promote plant resistance against pathogens and predators, and benefit human health (Aron & Kennedy, 2008; He & Giusti, 2010). Proanthocyanidins are synthesized by the sequential action of enzymes that are thought to form a weakly bound metabolon (Nakayama *et al*., 2019). This metabolon includes chalcone synthase (CHS), chalcone isomerase (CHI), flavanone 3-hydroxylase (F3H), flavonoid 3′-hydroxylase (F3′H), flavonoid 3′5′-hydroxylase (F3′5′H), dihydroflavonol 4-reductase (DFR), anthocyanin synthase (ANS), leucoanthocyanidin reductase (LAR), and anthocyanidin reductase (ANR) (Xie et al., 2003; Liu *et al*., 2013; Schaart *et al*., 2013). Of these enzymes, LAR and ANR are specific for proanthocyanidin biosynthesis.

The genes encoding these enzymes are regulated at the transcriptional level by members of various transcription factor (TF) families, including MYB, basic helix-loop-helix (bHLH), WRKY, WD40, zinc finger, and MADS box proteins (Grotewold, 2006; Petroni and Tonelli, 2011). These TFs act independently or in combination with cofactors to activate or repress flavonoid structural genes in a temporally and spatially controlled manner (Xu *et al*., 2014; Stracke *et al*., 2007; Xu *et al*, 2017). Among these TFs, MYB proteins have emerged as key players in the regulation of specific branches of flavonoid biosynthesis (Wang *et al*, 2017; Wang *et al*, 2019; Cao *et al*, 2019). MYB proteins have a conserved DNA binding MYB domain in the N-terminal, but their C termini are diverse and important for their transcriptional regulatory activity.

MYB proteins with an R2R3-type MYB domain (R2R3 MYBs) regulates flavonoid biosynthesis and are divided into different subgroups (SGs) regulating distinct biological functions (Dubos *et al*., 2010). R2R3 MYB proteins belonging to SG5 regulate proanthocyanidin biosynthesis (Nesi et al., 2001; Stracke et al., 2001). SG5 R2R3 MYBs contain a conserved basic bHLH-interacting motif ([DE]Lx(2)[RK]x(3)Lx(6)Lx(3)R) (Zimmermann *et al*., 2004) and form a trimeric MBW complex with one bHLH TF and one WD40 protein (Ramsay & Glover, 2005; Debeaujon *et al*., 2003; Zimmermann *et al*., 2004; Gonzalez *et al*., 2008). The regulation of proanthocyanidin biosynthesis is well characterized in Arabidopsis (*Arabidopsis thaliana*), where an MBW complex consisting of MYB123 (also named TRANSPARENT TESTA2 [TT2]), bHLH42 (also named TT8), and TRANSPARENT TESTA GLABRA1 (TTG1) regulates proanthocyanidin biosynthesis in seeds (Nesi *et al*., 2001; Baudry *et al*., 2004). Various TT2-like R2R3 MYB regulators have also been characterized in crops like barrel clover (*Medicago truncatula*) (*Mt*. PROANTHOCYANIDIN REGULATOR [MtPAR]; PA1-type MtMYB5 and MtMYB14; Verdier *et al*., 2012; Liu *et al*., 2014), grapevine (*Vitis vinifera*) (VvMYBPA1, VvMYBPA2, VvMYB5a, and VvMYBPAR; Deluc *et al*., 2006; 2008; Bogs *et al*., 2007; Terrier *et al*., 2009; Koyama *et al*. 2014), apple (*Malus domestica*) (MdMYB9, MdMYB11, and MdMYB12; Gesell *et al*., 2014; An *et al*., 2015; Wang *et al*., 2017), and peach (*Prunus persica*) (PpMYBPA1 and PpMYB7; Ravaglia *et al*., 2013; Zhou *et al*., 2015).

In contrast to MYB activators, involved in the regulation of flavonoid biosynthesis, work on MYB repressors has garnered increasing interest. MYB repressors are mainly categorized into R3 MYB and R2R2 MYBs, depending on the number MYB domain repeats they harbour. R3 MYB proteins have no known repressive amino acid motif and are thought to compete with R2R3 MYB activators for binding to their bHLH partners in the MBW complex (Guimil and Dunand, 2006; Zhang et al., 2009). Arabidopsis CAPRICE (CPC) and TRIPTYCHON (TRY) and petunia (*Petunia hybrida*) MYBx are prominent examples of known R3 MYB repressors (Zhu *et al*., 2009; Nemie-Feyissa *et al*., 2014). Arabidopsis MYB-LIKE2 (MYBL2), a negative regulator of anthocyanin biosynthesis (Dubos *et al*., 2008), also carries a single MYB domain but is more closely related to R2R3 MYBs and contains a TLLLFR amino acid repression motif in its C-terminal region (Matsui *et al*., 2008). R2R3 MYB repressors belong to SG4 and contain two conserved C-terminal amino acid motifs: C1 (lsrGIDPxT/NHR) and C2/EAR (pdLNLD/EL) (Aharoni et al., 2001; Dubos et al., 2010; Zhou et al., 2019). Other R2R3 MYB repressors have been identified: strawberry (*Fragaria* × *ananassa*) FaMYB1, *Fragaria chiloensis* FcMYB1 (Aharoni et al., 2001; Salvatierra *et al*., 2013), grapevine VvMYBC2-L1 (Huang *et al*., 2014), barrel clover MtMYB2 (Jun *et al*., 2015), peach PpMYB18 (Zhou *et al*., 2019), and poplar (*Populus trichocarpa*) PtMYB194 and PtMYB165 (Dawei Ma *et al*., 2018). As with R2R3 MYB repressors, R2R3 MYB activators are hypothesized to compete for binding to bHLHs, thus forming a transcriptional regulatory loop (Zhou *et al*., 2019).

Bananas (*Musa acuminata*) are a popular tropical and subtropical fruit crop cultivated in more than 120 countries (http://nhb.gov.in/report_files/banana/BANANA). The *Musa acuminata* reference genome sequence (D’Hont *et al*., 2012) provided a useful resource for functional genomics; however, little information is available about the transcriptional regulation of flavonoid biosynthesis in bananas. Here, we identified and characterized two R2R3-MYB activators (MaMYBPA) and four R2R2-MYB repressors (MaMYBPR) that regulate proanthocyanidin biosynthesis in bananas. We tested the role of *MaMYBPA* genes in proanthocyanidin regulation by ectopic expression in the Arabidopsis *tt2* mutant, deficient in proanthocyanidins. Transient overexpression of *MaMYB*s in banana fruits indicated that R2R3 MYB activators induce the expression of the genes encoding R2R3 MYB repressors, creating a negative feedback loop that modulates proanthocyanidin accumulation. Taken together, our results describe the molecular mechanisms underlying the transcriptional regulation of proanthocyanidin biosynthesis in *M. acuminata*.

## Material and Methods

### Plant Materials

Dessert banana, *Musa acuminata* (AAA genome) cultivar ‘Grand Nain’, was grown in the fields of the National Institute of Plant Genome Research (NIPGR), New Delhi, India. Roots, pseudostems, leaves, bracts, fruit peel, and pulp were collected and used for gene expression analysis. For transient experiments, fresh banana fruits were harvested from the field. The Arabidopsis Columbia-0 (Col-0) wild-type accession, the mutant *tt2-1* (NASC stock NW83, L*er* background), and transgenic Arabidopsis lines overexpressing *MaMYBPA* were grown under a 16-h-light/8-h-dark photoperiod at 22°C in a growth chamber (Percival AR-41L3). *Nicotiana benthamiana* plants were grown in a growth chamber with the same settings. The Arabidopsis cell suspension culture At7 (Trezzini *et al.,* 1993) is derived from the hypocotyl of the reference accession Columbia (Col) and was handled as described by Stracke *et al*. (2016).

### Multiple sequence alignment and phylogenetic analysis

The sequences of the R2R3-MYB domains were extracted and multiple sequence alignment of MaMYBPAs was carried out with the proteins listed in Supplemental Table S1. Alignment of each R2 and R3 domain was conducted using MAFFT v7 with default parameters (Katoh and Standley, 2013).

Phylogenetic analysis was carried out by aligning the R2R3-MYB domain sequences from banana, Arabidopsis, and selected plant landmark MYB sequences (Stracke, *et al*., 2014; Pucker *et al.,* 2020) using ClustalW with default parameters (Thompson *et al*., 1994). The maximum likelihood method was used with 1,000 bootstrap replicates using MEGA X software (Kumar *et al*., 2018). The tree was visualized with iTOLv6.4 (https://itol.embl.de/).

### *In silico* gene expression analysis

Banana RNA-Seq datasets were retrieved from the Sequence Read Archive (SRA, https://www.ncbi.nlm.nih.gov/sra) under BioProjects PRJNA182724, PRJNA79731, PRJNA316844, PRJNA316845, and SAMN00794551 for embryonic cell suspension (EC), seedling shoot (SS), seedling shoot (SR), and root (RT); leaf stages young (YL), adult (AL), and old leaf (OL); pulp stages S1–S4; and peel stages S1–S4. STAR v2.5.1b (Dobin *et al*., 2013) was applied to align the reads to the banana reference genome in 2-pass mode. Reads were considered mapped if the identity between read and the genome exceeded 95% and covered at least 90% of read length (Haak *et al*., 2018). Counts v1.5.0-p3 (Liao *et al*., 2014) was deployed with default settings to quantify gene expression. The count tables were processed and combined with previously developed Python scripts (Haak *et al*., 2018). Heatmap construction was performed with Hierarchical Clustering Explorer 3.5 and the hierarchical clustering of genes was executed according to the Euclidean distance method (Seo *et al*., 2006).

### Gene expression analysis

Total RNA was isolated from banana organs using a Spectrum Plant Total RNA Kit (Sigma Aldrich, India) and treated with RNAse-free DNase I (Fermentas Life Sciences). Total RNA was subjected to reverse transcription to generate first-strand cDNA using oligo(dT) primers (MBI Fermentas). The PCR mix contained 1 μL of diluted cDNA (corresponding to 10 ng total RNA), 5 μL of 2x SYBR Green PCR Master Mix (Applied Biosystems, USA), and 5 nM of each gene-specific primer (Table S2) in a final volume of 10 μL and transcript levels were measured on a 7500 Fast Real time PCR System (Applied Biosystems). Banana *Actin1* (GenBank No. AF246288) was used to normalize transcript abundance. Relative transcript levels were calculated using the cycle threshold (Ct) 2^−ΔΔCT^ method (Livak and Schmittgen, 2001). Three biological and three technical replicates were analysed for each transcript.

### Cloning of transcriptional regulators

The full-length coding sequences (CDSs, without the stop codon) of the MaMYB activator genes *MaMYBPA1* and *MaMYBPA2*, MaMYB repressor genes *MaMYBPR1* to *MaMYBPR4*, as well as *MaMYC2*, *MaMYC3*, *MaMYC7*, *MaMYC9*, *MaMYC13*, and *MaTTG1* (Ma04_35650) were amplified using first-strand cDNA of *M. acuminata* fruit tissue using a set of primers containing Gatewa attB sites designed on the basis of information from the Banana Genome Hub (Table S2). The resulting amplicons were cloned into the Gateway vector pDONRzeo (Invitrogen) and transformed into *E. coli* TOP10 cells. Sanger sequencing of entry clones was performed by the NIPGR sequencing core facility (New Delhi, India).

### Subcellular localization of transcriptional regulators

Entry clones harbouring the coding sequences of *MaMYBPA1*, *MaMYBPA2*, *MaMYBPR1*– *MaMYBPR4*, *MaMYC2*, and *MaMYC9* were recombined by Gateway LR reaction into the binary vector pGWB441 encoding the C terminus of YFP under the control of the CaMV 35S promoter (Nakagawa *et al*., 2007). The resulting plasmids were transformed into Agrobacterium (*Agrobacterium tumefaciens*) strain GV3101::pMP90 (Koncz and Schell, 1986). Agrobacteria harbouring each construct were resuspended along with an Agrobacterium colony carrying a nuclear marker (NLS-RFP) (Kumar *et al*., 2018) in freshly made infiltration medium (10 mM MgCl_2_, 10 mM MES-KOH, pH 5.7, 150 μM acetosyringone) and co-infiltrated into the abaxial surface of *N. benthamiana* leaves and kept at 22°C for 48 h. YFP and RFP fluorescence was observed under a Leica TCS SP5 confocal laser-scanning microscope (Leica Microsystems, Wetzlar, Germany) at 514 nm excitation, 527 nm emission and 558 nm excitation, 583 nm emission wavelength, respectively.

### Ultra-high-performance liquid chromatography (UHPLC) and Liquid chromatography– mass spectrometry (LC-MS)

Before analysis, all samples were filtered through a 0.22-µm PVDF syringe filter (Merck, Germany). The filtrate was collected in a clear-glass HPLC vial (Agilent Technologies). Separation for qualitative and quantitative analysis of monomeric and oligomeric proanthocyanidins was performed as described in the supplementary information. The LC-MS analysis was performed using an UPLC system (Exion LC Sciex) coupled to a triple quadrupole system (QTRAP6500+, ABSciex) using electrospray ionization. The voltage was set to 5,500 V for positive ionization with the MS parameters described in the in the supplementary information.

### Yeast two hybrid assay

For yeast two-hybrid (Y2H) assays, the *MaMYBPA*s, *MaMYBPR1*–*MaMYBPR4*, and *MaMYC* coding sequences were recombined into the prey vector pGADT7g (Clontech Laboratories Inc.) or the bait vector pGBKT7g (Clontech Laboratories Inc.). The resulting plasmids were co-transformed into Y2HGold (Clontech Laboratories Inc.) and cultured at 30°C on synthetic defined (SD) medium lacking Trp and Leu. For interaction tests, the yeast colonies carrying the appropriate bait and prey clones were restreaked onto medium lacking Trp, Leu, His, and Ade (QDO) with 40 mg/mL X-gal, 20 ng/mL aureobasidin (AbA), and 5 mM 3-amino-1,2,4-triazole (3-AT). p53/large T antigen interaction (Pipas and Levine, 2001) was used as positive control and empty pGADT7g/pGBKT7g vectors as a negative control.

### Bimolecular fluorescence complementation (BiFC) assay

The entry clones carrying the coding sequences of *MaMYBPA*s, *MaMYBPR1*–*MaMYBPR4*, and *MaMYC* were recombined into the vector 35S-pSITE-nYFP-C1 or 35S-pSITE-cYFP-N1 (Martin *et al*., 2009). The resulting plasmids were transformed individually into Agrobacterium strain GV3101::pMP90 (Koncz and Schell, 1986). Agrobacteria harbouring the constructs encoding the nYFP and cYFP fusion proteins to be tested were resuspended in freshly prepared infiltration medium (10 mM MgCl_2_, 10 mM MES-KOH, pH 5.7, and 150 μM acetosyringone) and co-infiltrated in 1:1 ratio into *N. benthamiana* leaves. Empty vectors (*nYFP*, *cYFP*) were used as a negative control. After incubation for 48 h at 22°C, YFP fluorescence was detected using a Leica TCS SP5 confocal laser-scanning microscope with 514-nm excitation and 527-nm emission wavelengths.

### Luciferase complementation imaging

The full-length coding sequences of *MaMYBPA1* and *MaMYBPA2* were cloned into the pCAMBIA1300-nLUC vector, while the entry clones of *MaMYC2*, *MaMYC7*, and *MaMYC9* were cloned into the vector pCAMBIA1300-cLUC (Chen et al., 2008). Agrobacterium strain GV3101 carrying the appropriate *nLUC* and *cLUC* constructs was individually resuspended in freshly prepared infiltration medium (10 mM MgCl_2_, 10 mM MES-KOH, pH 5.7, and 150 μ acetosyringone) and co-infiltrated in a 1:1 ratio into *N. benthamiana* leaves. Infiltrated leaves were infiltrated with 100 µM D-luciferin and LUC activity was quantified after 48 h using a low-light CCD imaging apparatus (Bio-Rad).

### Complementation analysis in Arabidopsis

Entry clones harbouring the *MaMYBPA* coding sequences were recombined into the binary destination vector pMDC32 (Curtis and Grossniklaus, 2003) carrying a double CaMV 35S promoter overexpression and hygromycin resistance cassette for selection of transgenic plants. The resulting plasmids were transformed into Agrobacterium strain GV3101::pM90RK (Koncz and Schell, 1986) by electroporation and then transformed into the proanthocyanidin-deficient Arabidopsis *tt2-1* mutant by the floral dip method (Clough and Bent, 1998). Selection of T_1_ transgenic plants was conducted on Murashige and Skoog (MS) solid medium containing 15 mg/L hygromycin. Resistant seedlings were transferred to soil and grown in a plant growth chamber to set seeds. T_3_ lines were analysed for proanthocyanidin accumulation.

### DMACA staining of seeds and determination of proanthocyanidin contents

4-Dimethylaminocinnamaldehyde (DMACA) staining was performed as previously described (Li *et al*., 1996; Wang *et al*., 2017). The proanthocyanidin content was measured with DMACA reagent as previously described (Pang *et al.,* 2007). For proanthocyanidin quantification, catechin was used to prepare a calibration curve (200, 400, 600, 800, and 1,000 µg/mL) and contents were determined as catechin equivalents. Samples, blanks, and standards were read within 15 min of staining on a FLUOstar Omega Microplate Reader (BMG, Germany) with a 640-nm emission filter.

### Structural analysis of the *MaANR*, *MaANS*, and *MaLAR* promoters

The 1234-, 1518, and 1516-bp sequences upstream of the translation start of the *MaANS*, *MaLAR*, and *MaANR* genes, respectively, were analysed for the presence of putative MYB and bHLH binding sites. Several sites were identified in the NEW PLACE database (http://www.dna.affrc.go.jp/PLACE/signalscan.html) (Higo *et al.,* 1999) and from previously identified motifs (Hartmann *et al.,* 2005; Urao *et al.,* 1993; Solano *et al.,* 1995 and Wang *et al.,* 1997).

### Yeast one-hybrid (Y1H) assay

The Matchmaker Gold Yeast 1-hybrid system (Clontech) with the Y1H gold yeast strain was used. Promoter fragments of *MaANR* (*proMaANR-938*), *MaANS* (*proMaANS-756*), and *MaLAR* (*proMaLAR-839*) were amplified using a set of primers with Gateway attB sites (supplementary Table S2) and cloned into the pABAi vector (Cat. No. 630491, Takara Bio USA, Inc.). The *MaMYBPA1* and *MaMYBPA2* coding sequences were cloned into the pGADT7-GW vector (Clontech) to generate effector constructs. The pABAi plasmids carrying the promoter fragments were linearised and transformed into Y1H Gold strain according to the Matchmaker manual. Effector plasmids and empty pGADT7-GW vector control were transformed into the Y1H strains with genome-integrated reporter constructs. To test for interaction, the yeast colonies containing effector and reporter constructs were restreaked onto medium containing AbA. Yeast growth was checked under different concentrations of AbA to determine the minimum inhibitory concentration (MIC) values and check possible interaction. The MIC values for *proMaANR* and *proMaANS* were 900 ng/mL AbA and 950 ng/mL for *proMaLAR*.

### Co-transfection in Arabidopsis At7 protoplasts

The entry clones of the promoter fragments of *MaANS-1234*, *MaLAR-1518*, and *MaANR-1516* genes were amplified using primers containing Gateway attB sites (Supplementary Table S2). To generate effector constructs overexpressing transcription factor genes from the CaMV 35S promoter, the *MaMYB*s, *MaMYC*s, and *MaTTG1* coding sequences were recombined into the vector pBTdest (Baudry *et al*., 2004). The cultivation of At7 cells, co-transfection of effector and reporter constructs into protoplasts, and the determination of activation capacity were performed as described by Stracke *et al*. (2016).

### Dual-luciferase reporter assay

To test the transactivation potential of the regulators MaMYBPA1, MaMYBPA2, and MaMYBPR1-MaMYBPR4, the entry clones of the above-mentioned promoters (proMaANS-1234, proMaLAR-1518, and proMaANR-1516) were recombined into p635nRRF containing 35S:REN. The effector constructs harbouring *MaMYBPA1*, *MaMYBPA2*, *MaMYBPR1*-*MaMYBPR4*, and *MaMYC*s were recombined into the pBTdest vector. The reporters were co-infiltrated with different effectors into *N. benthamiana* leaves. Firefly luciferase and REN activity were measured after 48 h in extracts from infiltrated leaf discs using the Dual-Luciferase Reporter Assay System (Promega, USA) according to the manufacturer’s protocol. The samples were quantified using the POLARstar Omega multimode plate reader (BMG Labtech, Germany), and firefly luciferase activity was normalised to REN activity.

### Transient overexpression in immature banana fruit slices

For transient overexpression studies, the coding sequences of *MaMYBPA1*, *MaMYBPA2*, *MaMYBPR1* to *MaMYBPR4*, and *MaMYC* were recombined into the vector pANIC6B (Mann *et. al*., 2012) and placed under the control of the constitutive maize *Ubiquitin1* (*ZmUBI1*) promoter. The T-DNA of pANIC6B additionally contains a *proPvUBI1-GUSplus-nosT* expression cassette to follow successful transformation by GUS activity. The plasmids were transformed into Agrobacterium strain GV3101::pMP90 (Koncz and Schell, 1986), and transient overexpression experiments were performed according to Matsumoto *et al*. (2009) with minor modifications. In brief, Agrobacterium cells carrying the plasmid were resuspended in infiltration medium (one-tenth strength MS salts, one-tenth strength B5 vitamins, 20 mM MES-KOH, pH 5.7, 2% [w/v] sucrose, 1% [w/v] glucose, and 200 μM acetosyringone) and kept in the dark for 3 h. Fruit discs were soaked in the Agrobacterium suspension and vacuum-infiltrated for 15 min. Excess Agrobacterium-containing medium was wiped from fruit slices, and transformed fruit discs were kept on cultivation medium (same as infiltration medium) for 3 d and then used for GUS staining (transformation efficiency), DMACA staining (detecting proanthocyanidins), proanthocyanidin quantification, and gene expression analysis.

## Results

### Identification of proanthocyanidin activating R2R3 MYB regulators from *M. acuminata*

In an earlier study, *in silico* analysis identified putative R2R3 MYB regulators in *M. acuminata (*Pucker *et al.,* 2020). We focused here on two homologous R2R3 MYB activators (Ma03_p07840 and Ma10_p17650) with high similarity to known proanthocyanidin regulators and named them *MaMYBPA1* (Ma03_g07840) and *MaMYBPA2* (Ma10_g17650), with PA standing for proanthocyanidin activator. Previous reports suggested that along with activators, repressors also act in the flavonoid pathway. We identified four candidate R2R3 MYB proanthocyanidin repressors (PR) designated MaMYBPR1, MaMYBPR2, MaMYBPR3, and MaMYBPR4 all having conserved C1 (lsrGIDPxT/NHR), C2 (pdLNLD/EL), and TLLLFR repressor motifs at their C termini (Figure S1a).

We explored the phylogenetic relationship of MaMYBPA and MaMYBPR proteins with known R2R3 MYB activators and repressors from monocot and dicot plants. The four MaMYBPRs form a clade with previously known proanthocyanidin-specific repressors, suggesting a role as flavonoid-specific repressors (Figure 1a). Similarly, MaMYBPA1 and MaMYBPA2 formed a clade with known R2R3 MYB activators like Arabidopsis TT2, strawberry FaMYB9, apple MdMYB9, and homologues from fruit crops like grapevine (Figure 1a). Multiple sequence alignment of MaMYBPA1 and MaMYBPA2 with previously characterized proanthocyanidin regulators from Arabidopsis (AtTT2), wild apple (*Malus sieversii*; MdMYB9, MdMYB11, and MdMYB12), strawberry (FaMYB9), and grapevine (VvMYBPA2) demonstrated a high degree of conservation between their N-terminal R2R3 MYB domain and the bHLH-interacting consensus motif [D/E]Lx2[R/K]x3Lx6Lx3R, while the C-terminal region contained the SG5 motif (IRTKA[I/L]RC) (Zhao *et al*., 2013) was specific for R2R3 MYBs that positively regulate proanthocyanidin biosynthesis (Figure 1b).

**Figure 1.**
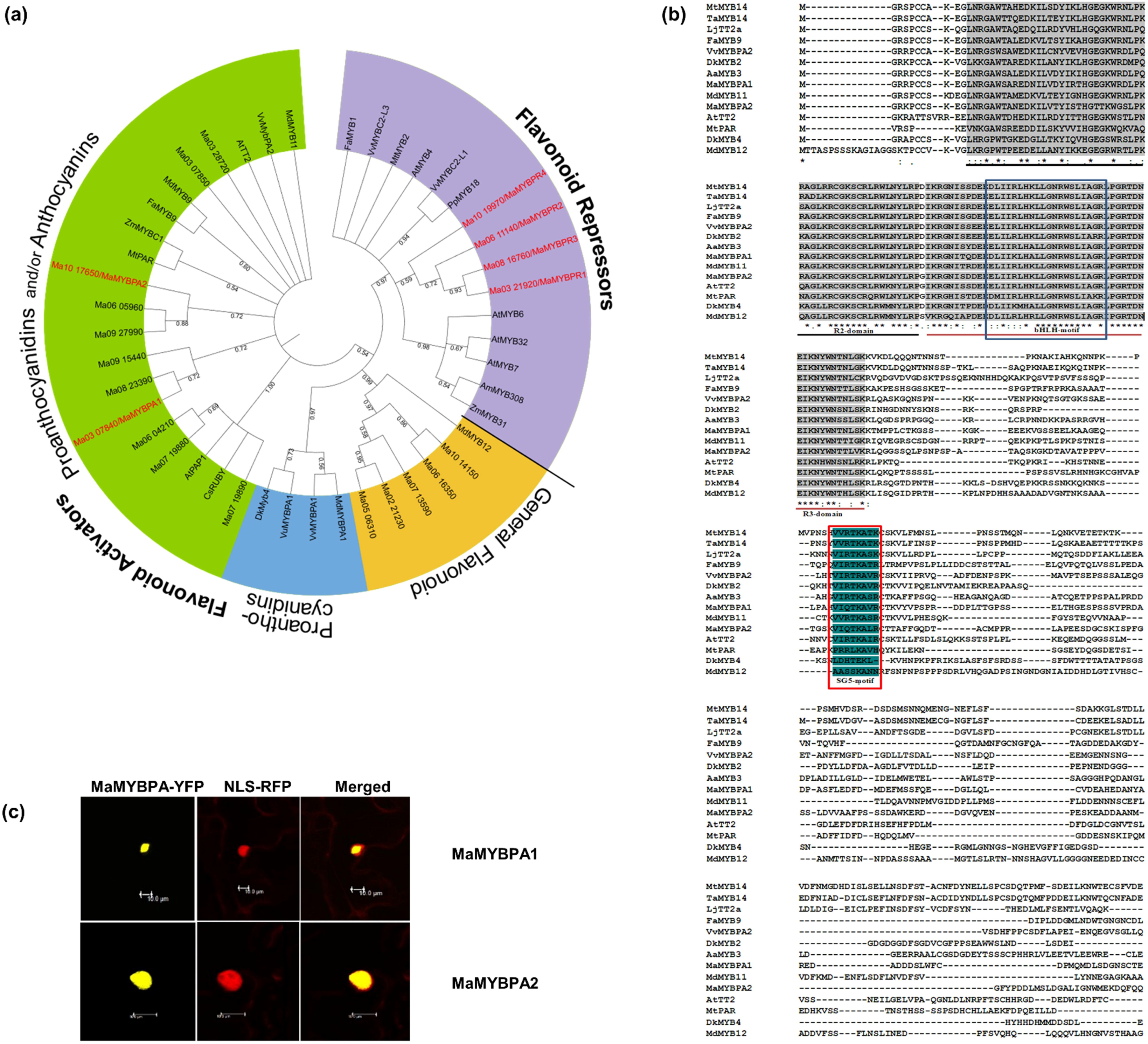
Candidate proanthocyanidin specific R2R3 MYB regulators from *Musa acuminata*: phylogenetic analysis and subcellular localization. (a) Phylogenetic tree of R2R3-MYB activators and repressors, constructed using the Maximum-likelihood method in MEGA X software and visualized with iTOLv6.4 (https://itol.embl.de/). Bootstrap values are shown with numbers on each branch. Numbers on each branch indicate branch length. Proanthocyanidin-specific regulators from *M. acuminata* were identified based on their clustering with known flavonoid-specific R2R3 MYBs and are highlighted in red colour. (b) Amino acid sequence alignment of the characterized proanthocyanidin-specific R2R3 MYB regulators (MaMYBPA1 and MaMYBA2) from *M. acuminata* with known proanthocyanidin-specific R2R3-MYB regulators from other plant species. The R2 and R3 repeats of the MYB domain, the bHLH interaction motif, and the subgroup 5 (SG5) defining motif are indicated. **(c)** The MaMYBPA-YFP fusion protein localizes to the nucleus, as evidenced by colocalization with the nuclear marker NLS-RFP, in Agrobacterium-infiltrated *N. benthamiana* leaves analysed by confocal microscopy. Representative images are shown. Scale bars = 10 µm.

We determined the subcellular localization of MaMYBPA1 and MaMYBPA2 by transiently co-infiltrating constructs encoding a yellow fluorescent protein (YFP) fusion in *Nicotiana benthamiana* epidermal cells and the nuclear marker NLS-RFP (red fluorescent protein [RFP] fused to a nuclear localization signal [NLS]). We detected YFP fluorescence exclusively in the nucleus strictly colocalized NLS-RFP (Figure 1c), indicating that MaMYBPA1 and MaMYBPA2 are nuclear proteins. We obtained similar results with MaMYBPRs, in agreement with their predicted function as transcriptional repressors (Figure S1b).

### *MaMYBPA* expression patterns correlate with those of their structural target genes and metabolite contents

We used a transcriptome deep sequencing (RNA-Seq) dataset comprising banana samples representing different organs and developmental stages: embryonic cell suspension (EC), seedling shoot (SS), seedling root (SR), root (R), young (YL), adult (AL), and old leaf (OL) stages, pulp stages S1–S4, and peel stages S1–S4 to analyse the expression of *MaMYBPA1* and *MaMYBPA2* genes and of three proanthocyanidin biosynthesis genes *MaANS*, *MaLAR*, and *MaANR*. *MaMYBPA1* and *MaMYBPA2* were highly expressed during the early stage of fruit pulp (S1) (Figure 2a). *MaANS* and *MaANR* were also highly expressed during the S1 stage of pulp development (Figure 2a).

**Figure 2.**
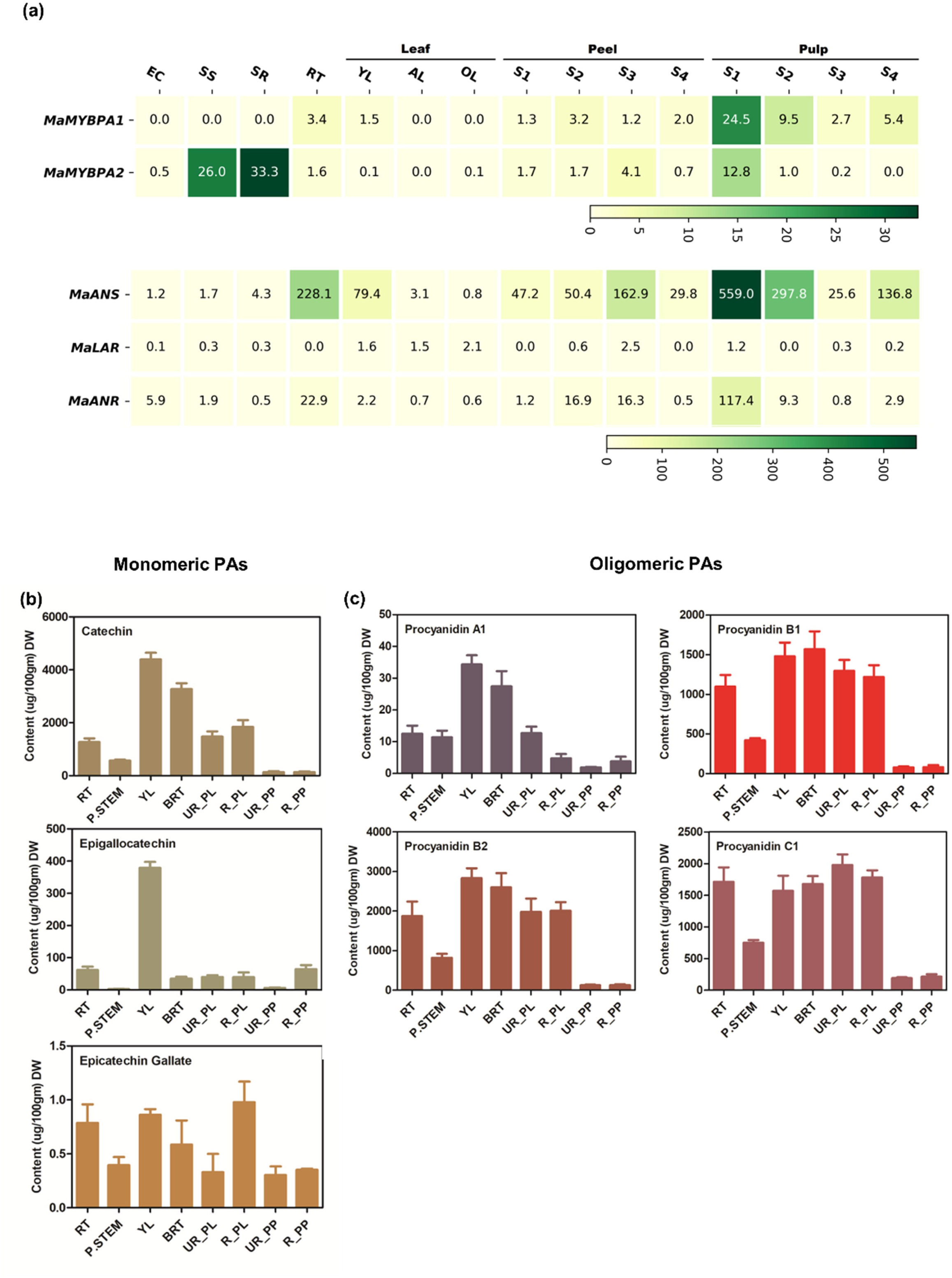
Proanthocyanidin-related gene expression and metabolite contents in different organs of *M. acuminata*. (a) Heatmap representation of the expression of *MaMYBPA*s and candidate PA-branch biosynthesis genes (*MaANS*, *MaANR*, and *MaLAR*) in embryogenic cells (EC), seedling shoots (SS), seedling roots (SR), roots (RT), and different developmental stages of leaves (young, adult, and old), peel (stage 1 to stage 4 [S1 to S4]), and pulp (stage 1 to stage 4 [S1 to S4]) organs/tissues. FPKM values for the indicated genes were extracted from publicly available RNA-seq datasets and plotted according to the colour scale. *ANS*, *Anthocyanidin synthase*; *ANR*, *Anthocyanidin reductase*; *LAR*, *Leucoanthocyanidin reductase*. (b, c) Contents for monomeric PAs (b) and oligomeric PAs (c) in different *M. acuminata* organs. Compounds were quantified by separating methanolic extracts using UHPLC coupled with MS. Data are shown as means ± standard deviation (SD) of two independent biological replicates each having three technical replicates.

We also conducted a targeted metabolite profiling using ultra-high-performance liquid chromatography (UHPLC) and liquid chromatography–mass spectrometry (LC-MS) to quantify three monomeric and four oligomeric proanthocyanidin forms in different organs (Figure 2b, 2c; Figure S2a and S2b). This showed that monomeric and oligomeric forms are abundant in young leaves, bracts, and peels of banana fruits (Figure 2c). In contrast to the expression data above, we did not observe a positive correlation between gene expression profiles and metabolite contents.

### MaMYBPA1 and MaMYBPA2 interact with MaMYC

As proanthocyanidin biosynthesis relies on a regulatory MBW complex, we sought to identify partner bHLH and WD40 proteins. Phylogenetic analysis revealed five ENHANCER OF GLABRA3 (EGL3)-like and eight TT8-like bHLH proteins encoded by the banana reference genome. We named the bHLH proteins MaMYC1 to MaMYC13, ordered based on the chromosomal locations (Figure S3). We also conducted a phylogenetic analysis of MaTTG1, which yielded two TTG1-like proteins named MaTTG1.1 and MaTTG1.2 (Figure S3). As MaMYBPA1 and MaMYBPA2, both contain a bHLH-binding motif ([DE]Lx2[RK]x3Lx6Lx3R), we tested whether they are able to interact with the MaMYCs. We determined that MaMYBPA1 interacts with MaMYC2 and MaMYC7 in yeast two-hybrid (Y2H) assays, while MaMYBPA2 interacted only with MaMYC9 (Figure 3a), supporting a direct interaction between the two R2R3 MYBs and MaMYC proteins. We also performed bimolecular fluorescence complementation (BiFC) and luciferase complementation imaging (LCI) assays in *N. benthamiana* leaves to extend these results *in planta*. Indeed, we detected the reconstitution of YFP, indicative of protein–protein interaction, in BiFC assays when constructs encoding MaMYBPA1-YFP^N^ and MaMYC2-YFP^C^, MaMYBPA1-YFP^N^ and MaMYC7-YFP^C^, and MaMYBPA2-YFP^N^ and MaMYC9-YFP^C^ were transiently co-infiltrated into *N. benthamiana* leaves (Figure 3b). We obtained comparable results via LCI assays demonstrating an interaction between MaMYBPA1 and MaMYC2 and MaMYC7, as well as between MaMYBPA2 and MaMYC9 (Figure 3c). We also tested the localisation of MaMYC2 and MaMYC9 and found that both MaMYB-MaMYC co-localized in the nucleus (Figure S4).

**Figure 3.**
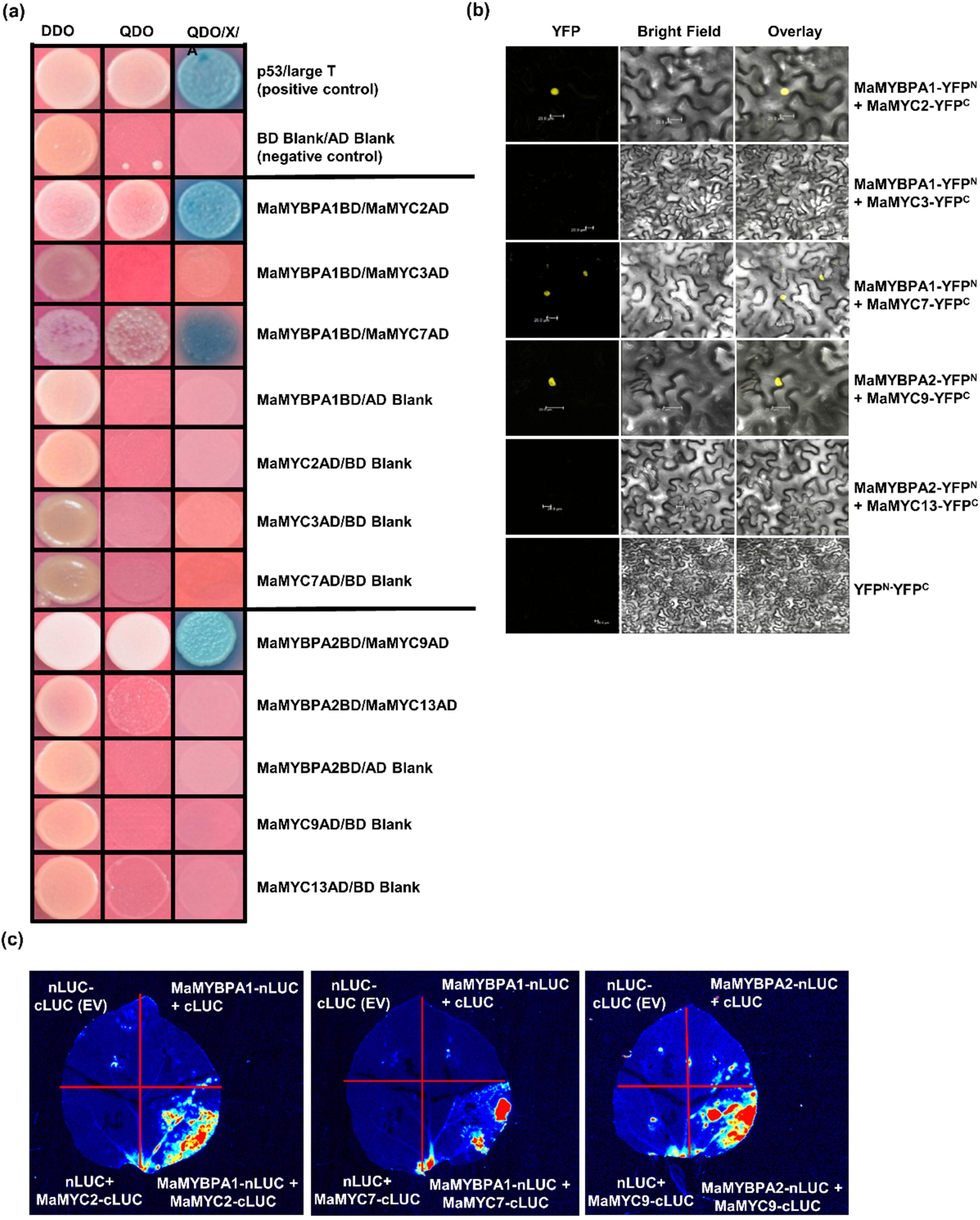
Interaction between MaMYBPA and MaMYC proteins. (a) Yeast two-hybrid assays showing the interaction of MaMYBPA1 with MaMYC2 and MaMYC7, and of MaMYBPA2 with MaMYC9. AD, GAL4 activation domain; BD, GAL4 DNA-binding domain; DDO, double dropout synthetic defined (SD)–Leu–Trp medium; QDO, quadruple dropout SD–Leu–Trp–Ade–His medium; QDO/X/A, QDO +X-α-Gal + AbA medium. (b) Bimolecular fluorescence complementation (BiFC) assay indicating the *in planta* interaction of MaMYBPA1 with MaMYC2 and MaMYC7 and MaMYBPA2 with MaMYC9. *YFP^N^* + *YFP^C^* constructs were used as a negative control. Scale bars = 20 µm. (c) Luciferase complementation imaging (LCI) assay showing the interaction between MaMYBPA1 and MaMYC2 and MaMYC7, and of MaMYBPA2 with MaMYC9. The following three pairs of constructs were used as negative controls: *MaMYBPA-nLUC+cLUC*, *nLUC+MaMYC-cLUC*, and *nLUC+cLUC*.

### *MaMYBPA1* and *MaMYBPA2* partially rescue the proanthocyanidin-deficient phenotype of the Arabidopsis *tt2-1* mutant

We overexpressed *MaMYBPA1* and *MaMYBPA2* in the Arabidopsis *tt2-1* mutant, which is defective in the proanthocyanidin regulatory R2R3 MYB TT2 and has yellow seeds rather than the normal brown seeds. We stained the seeds of several stable transgenic lines harbouring the *35Spro:MaMYBPA1* or *35Spro:MaMYBPA2* transgenes with 4-dimethylaminocinnamaldehyde (DMACA), which revealed a rescue of proanthocyanidin accumulation, as evidenced by the blue-black colour of the stained seeds (Figure 4a). Furthermore, spectrophotometric quantification indicated that transgenic lines L-1, L-7, and L-8 carrying the *35Spro:MaMYBPA1* transgene and lines L-4, L-10, and L-13 harbouring *35Spro:MaMYBPA2* accumulate proanthocyanidins in their seeds, whereas *tt2-1* mutant seeds lacked detectable proanthocyanidins. Notably, none of the transgenic lines restored proanthocyanidin contents to wild-type levels (Figure 4b). Nevertheless, we concluded that MaMYBPA1 and MaMYBPA2 are likely proanthocyanidin regulators in banana.

**Figure 4.**
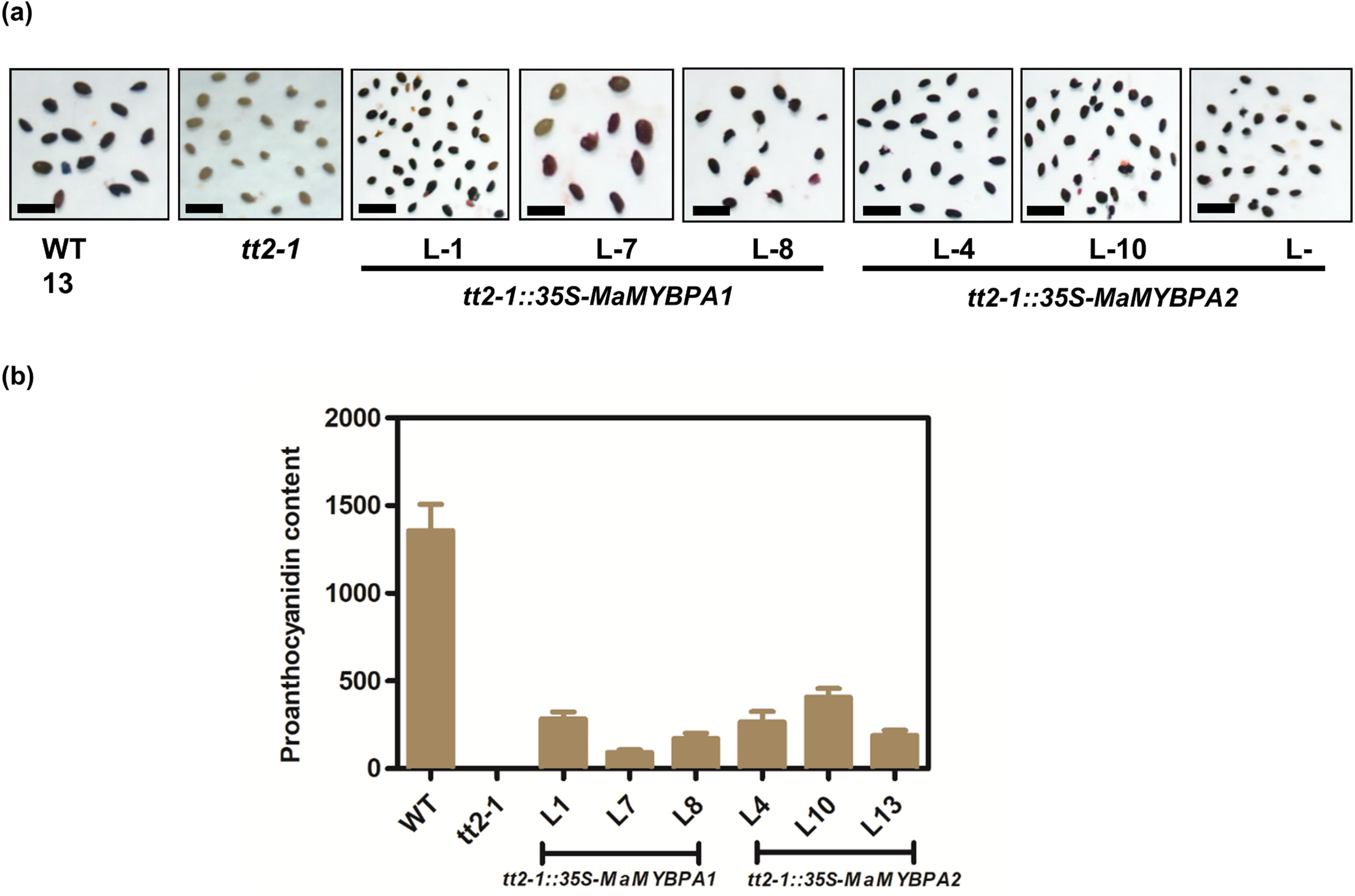
*MaMYBPA* overexpression partially rescues the Arabidopsis *tt2-1* mutant phenotype. (a) DMACA-stained seeds of Arabidopsis L*er* wild type (WT), *tt2-1* loss-of-function mutant, and selected, independent transgenic *tt2-1 35Spro:MaMYBPA* transgenic lines. Scale bars = 5 µm. (b) PA contents of seeds. (c) Relative *LDOX* and *BAN* transcript levels from the seeds of Arabidopsis lines, as determined by RT-qPCR. Data are shown as means ± SD values of three biological and three technical replicates. *MaACTIN* was used as reference.

### MaMYBPAs bind to the *MaANS*, *MaANR*, and *MaLAR* promoters

To link TFs to their targets, we looked for putative MYB binding sites (MBS) and bHLH binding sites (BBS) in the promoters of structural proanthocyanidin biosynthesis genes. We identified five putative MBSs and six BBSs within a 1,234-bp *MaANS* promoter fragment (*proMaANS-1234*, upstream of the transcription start site). Likewise, we detected five MBSs and six BBSs sites in *MaLAR-1503* promoter (a *1234*-bp fragment upstream of *MaANR*), while *MaANR-1516* promoter contained six MBSs and five BBSs sites (a 1,516-bp fragment upstream of *MaANR*). Details of the putative binding sites are depicted in Figure 5a.

**Figure 5.**
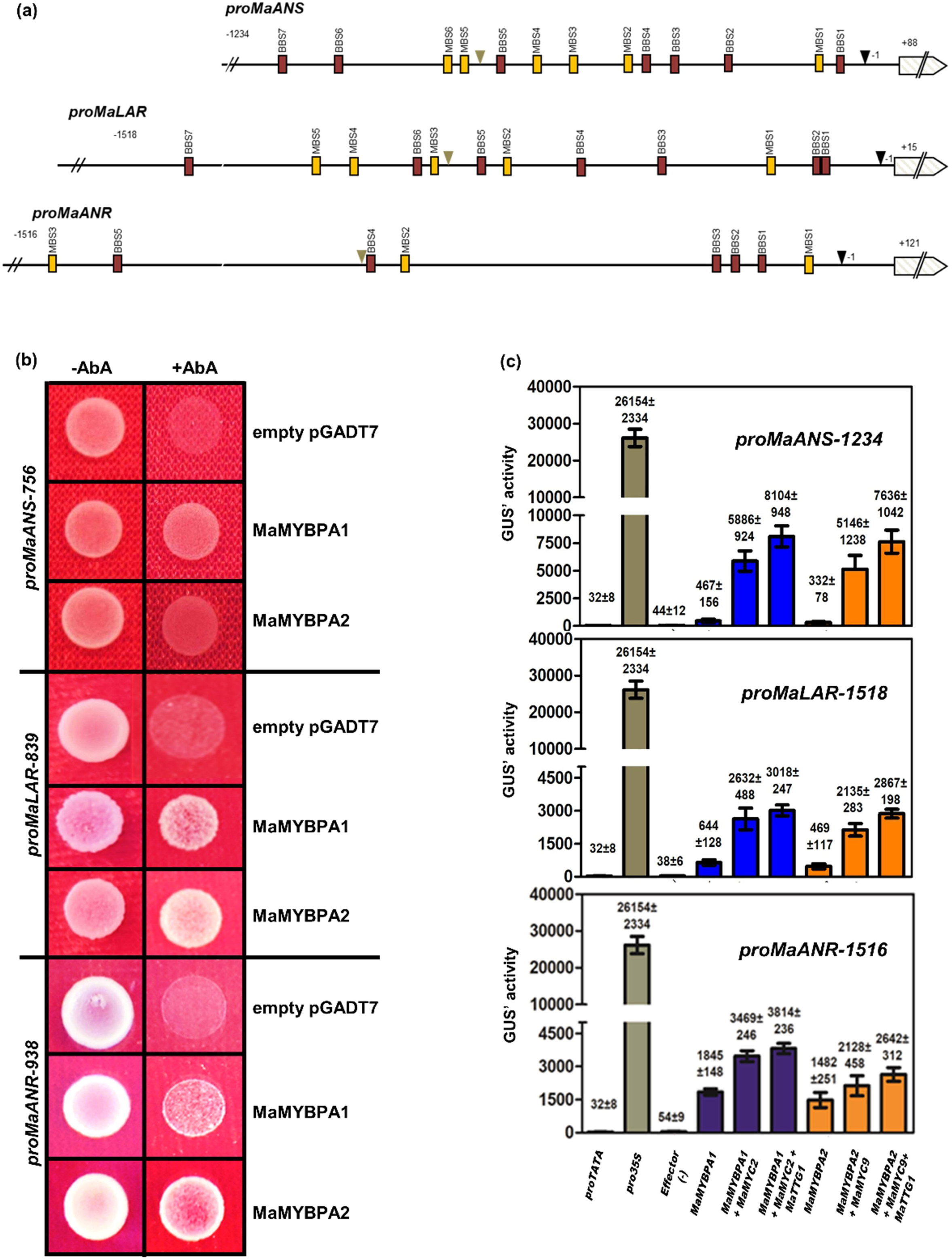
MaMYBPA proteins function in an MBW complex and can activate the promoters of PA biosynthesis genes. (a) Schematic diagram of the *MaANS*, *MaLAR*, and *MaANR* promoters. Putative *cis*-acting elements are indicated: MBS (yellow), MYB binding sites; G-box elements (red), bHLH binding sites (BBS). The black triangles mark the –1 position relative to the transcription start site (TSS). Positions +88, +121, and +15 show the translation start sites of *proMaANS*, *proMaANR*, and *proMaLAR,* respectively. Grey triangles indicate the length of the promoter fragments used for yeast one-hybrid (Y1H) analysis. (b) Results of Y1H experiments, testing the interaction between MaMYBPA1, MaMYBPA2, and *proMaANR*, *proMaANS*, and *proMaLAR.* Empty pGADT7 vector was used as negative control. (c) Results from co-transfection experiments in Arabidopsis protoplasts. Reporter constructs consisting of *GUS* driven by a 1234-bp *MaANS*, 1518-bp *MaLAR* and 1516-bp *MaANR* promoter fragment (reporter) were assayed for their transactivation by the 35S promoter-driven effectors *MaMYBPA1*, *MaMYBPA2*, *MaMYC2*, *MaMYC9*, and *MaTTG1*, either alone or in the indicated combinations. Data are shown as means of normalised GUS activity. Data from a set of six technical replicates are presented.

We then performed a yeast one-hybrid (Y1H) assay, which showed that MaMYBPA1 and MaMYBPA2 bind to *proMaANS-756*, *proMaLAR-839* and *proMaANR-938*, while the corresponding empty vector controls did not. These results indicated that *MaANS*, *MaANR*, and *MaLAR* are likely target genes of MaMYBPA1 and MaMYBPA2 (Figure 5b).

### MaMYBPA-containing MBW complexes activate the *MaANS*, *MaANR*, and *MaLAR* promoters

To assess the transactivation potential of MaMYBPA1 and MaMYBPA2 on the promoters of their target genes *proMaANS-1234*, *proMaANR-1516*, and *proMaLAR-1518 in planta*, we transiently transfected Arabidopsis protoplasts for a transactivation assay. To this end, we placed the β*-glucuronidase* (*GUS*) reporter gene under the control of each promoter and co-transfected the reporter with effector constructs, representing one or more TFs. Each MaMYBPA alone transactivated the *MaANS*, *MaANR*, and *MaLAR* promoters to some extent. However, co-transfection of MaMYBPA1 or MaMYBPA2 with MaMYC or MaMYC and MaTTG1 further induced the transactivation of the tested promoters (Figure 5c). These results indicated that MaMYBPA1 and MaMYBPA2 form functional MBW complexes with MaMYC and MaTTG1 *in planta* and transactivate the promoters of the proanthocyanidin biosynthesis genes *MaANS*, *MaLAR* and *MaANR*.

### A MaMYBPR-containing MBW complex binds to the *MaANS*, *MaANR*, and *MaLAR* promoters and represses their transactivation

To test the effect of the putative repressors on transactivation of the target gene promoters *proMaANS-1234*, *proMaLAR-1518*, and *proMaANR-1516*, we performed a dual-luciferase assay in *N. benthamiana* leaves. Accordingly, we placed the firefly luciferase (*LUC*) reporter gene under the control of each promoter fragment and transiently infiltrated each reporter construct with one or more effector construct (Figure 6a). In agreement with the transactivation assay in Arabidopsis protoplasts, we observed a strong increase in luminescence when *MaMYBPA1* or *MaMYBPA2* was overexpressed compared to the promoter constructs without any effector (effector-), suggesting that both TFs bind to *proMaANS*, *proMaLAR*, or *proMaANR* on their own (Figure 6b). However, the co-infiltration of *MaMYBPA1* or *MaMYBPA2* with *MaMYCs* transactivated the tested promoters to higher levels than with *MaMYBPA1* or *MaMYBPA2* alone. Importantly, the co-infiltration of *MaMYBPA1* or *MaMYBPA2* with *MaMYC* and effector constructs for the putative MaMYB repressors MaMYBPR2, MaMYBPR3, or MaMYBPR4 repressed promoter transactivation, as evidenced by the lower relative LUC activity measured (Figure 6b). These results suggested that MaMYBPRs act as repressors in the MBW complex.

**Figure 6.**
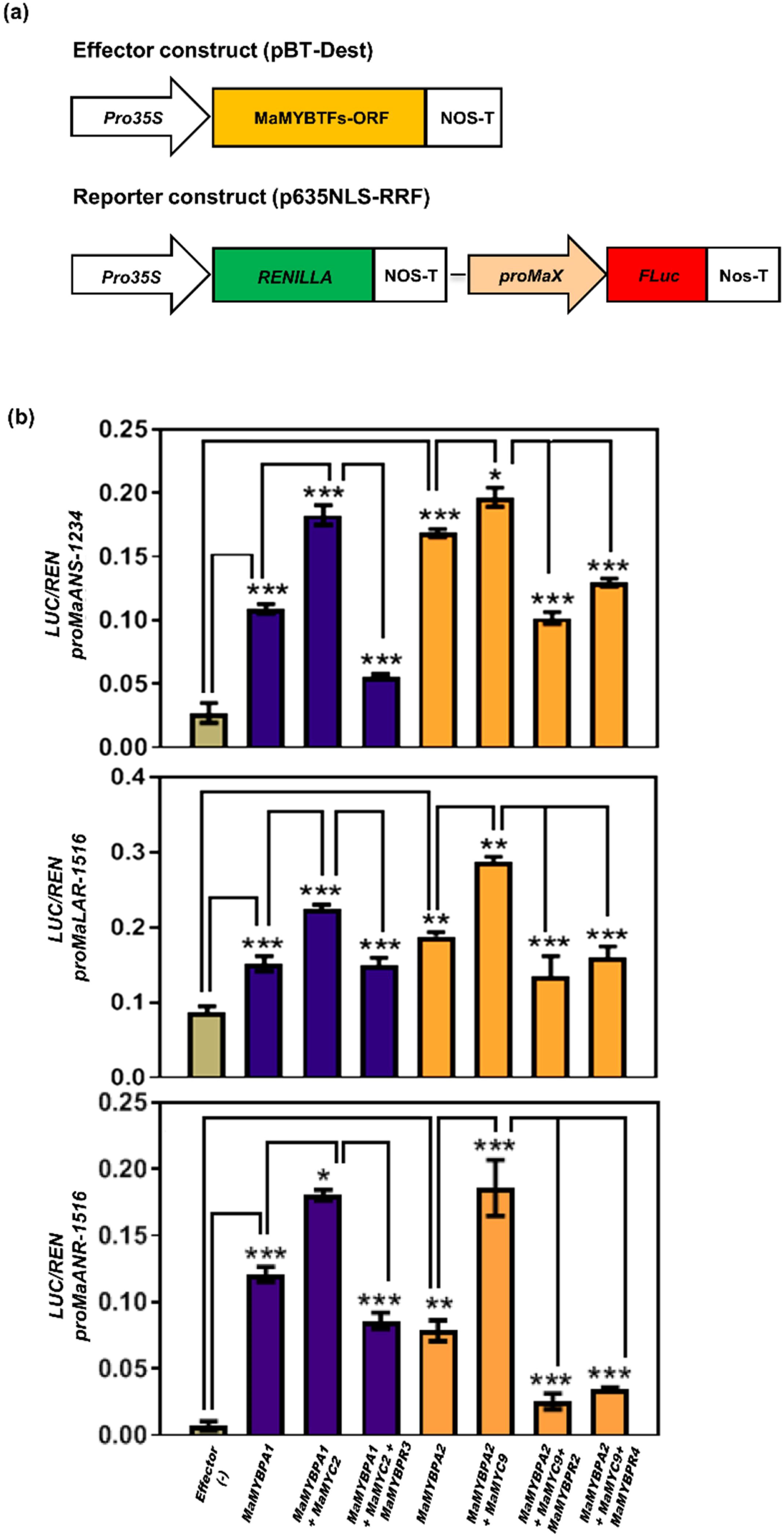
MaMYBPR-containing MBW complex binds to and represses the *MaANS*, *MaANR*, and *MaLAR* promoters. (a) Schematic diagram of the effector and reporter constructs used in the dual luciferase experiment. (b) Results from transient luciferase assays in *N. benthamiana* leaves. Constructs harbouring the firefly luciferase reporter gene (*LUC*) driven by 1234-bp *MaANS*, 1518-bp *MaLAR* and 1516-bp *MaANR* promoter fragments were transiently co-infiltrated with *MaMYBPA1*, *MaMYBPA2*, *MaMYBPR2-MaMYBPR4*, *MaMYC2*, and *MaMYC9* effector constructs, either alone or in the indicated combinations. *, *p* ≤ 0.05; **, *p* ≤ 0.01, ***, *p* ≤ 0.001, as determined by one-way ANOVA. Data are shown as means ± SD of four biological replicates.

### Transient overexpression of *MaMYBPA*s enhances proanthocyanidin accumulation in unripe banana fruits

We next transiently overexpressed *MaMYBPA1* and *MaMYBPA2* individually or together with their interacting bHLH partners (*MaMYBPA1*+*MaMYC2* and *MaMYBPA2*+*MaMYC9*) in unripe banana fruit slices. Each TF was driven by the maize *Ubiquitin1* (*ZmUBI1*) promoter, while the *GUS* reporter was driven by the *UBI1* promoter from *Phaseolus vulgaris* (*proPvUBI1*) as control for transient transformation (Figure 7a). We observed the successful transformation of fruit slices, as shown by GUS activity (Figure 7b). DMACA staining revealed the accumulation of proanthocyanidins in the transformed fruit slices. While we detected little accumulation of proanthocyanidins in the untransformed control, the amount of proanthocyanidins increased substantially in fruit slices transformed with *MaMYBPA* alone as well as in those co-transformed with *MaMYBPA* and *MaMYC* (Figure 7c).

**Figure 7.**
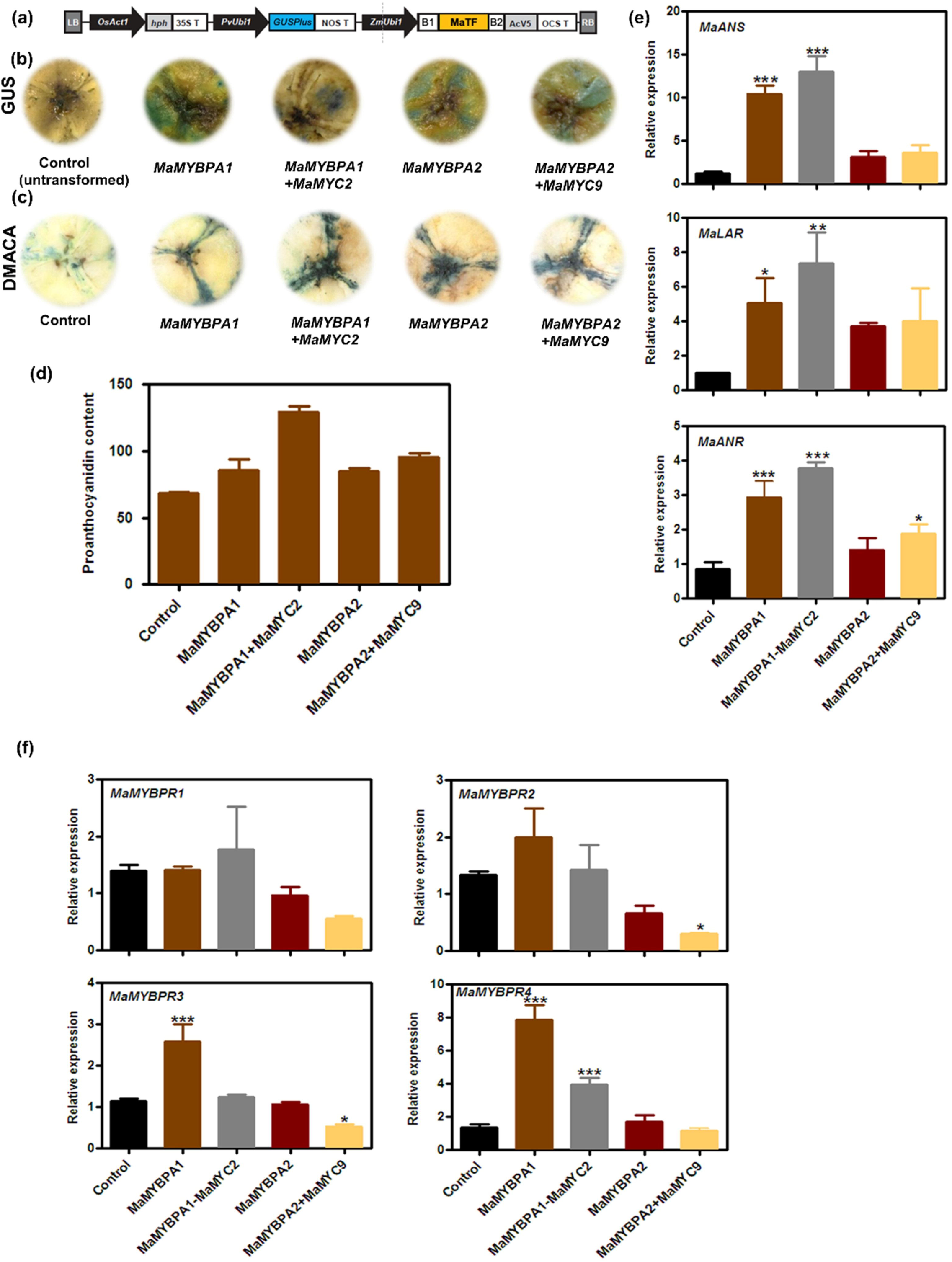
Functional analysis of *MaMYBPA1* and *MaMYBPA2* genes in banana fruits. (a) Schematic diagram of the reporter construct transiently expressed in unripe banana fruit slices via Agrobacteria infiltration. The *ZmUBI* promoter was used to drive the overexpression of *MaMYBPA1*, *MaMYBPA1+MaMYC2* (co-expression), *MaMYBPA2*, or *MaMYBPA2+MaMYC9* (co-expression), together with the *GUS* reporter construct driven by the pea (*Phaseolus vulgaris*) *Ubi1* promoter as an indicator of transient transformation. (b-d) GUS staining (b), DMACA staining (c), and PA contents (d) in transiently transformed fruit slices and control (empty vector). (e) Relative *MaANS*, *MaANR*, and *MaLAR* transcript levels in transiently transformed fruit slices and the control (empty vector), as determined by RT-qPCR. (f) Relative transcript levels for the four candidate R2R3-MYB repressor genes (*MaMYBL1–MaMYBL4*) in transiently transformed fruit slices and control (empty vector). Data are shown as means ±SD of three biological replicates each having three technical replicates. *, *p* ≤ 0.05; **, *p* ≤ 0.01, ***, *p* ≤ 0.001, as determined by one-way ANOVA.

We also quantified proanthocyanidins spectrophotometrically in transformed fruit slices, which confirmed the accumulation of the pigment in fruit slices transiently overexpressing each *MYB* alone or in combination with their MYC partner (Figure 7d). The overexpression of *MaMYBPA1* or *MaMYBPA2* alone resulted in comparable proanthocyanidin contents (Figure 7c), suggesting that the expression of *MaMYBPA1* and *MaMYBPA2* provides sufficient flux for the biosynthesis of proanthocyanidins in banana fruit pulp. Moreover, the combined overexpression of *MaMYBPA1* and *MaMYC2* or *MaMYBPA2* and *MaMYC9* further enhanced proanthocyanidin biosynthesis with highest proanthocyanidin accumulation in *MaMYBPA1* and *MaMYC2*. The expression levels of target structural genes showed a strong positive correlation with the proanthocyanidin profiles, as determined by RT-qPCR analysis. Indeed, relative *MaANS*, *MaLAR* and *MaANR*, transcript levels were induced to higher levels in fruits transiently co-expressing *MaMYBPA1*+*MaMYC2* compared to fruits expressing only *MaMYBPA1* or *MaMYBPA2*. Notably, the co-overexpression of *MaMYBPA2* and *MaMYC9* resulted in transcript levels similar to those with *MaMYBPA2* alone (Figure 7e).

### *R2R3 MYBPR*s are induced in *MaMYBPA-*overexpressing banana fruit slices

To dissect the effects of MaMYBPA and MaMYC on the transcript levels of the repressor-type R2R3 MYB genes, we performed RT-qPCR in banana fruit slices transiently overexpressing the *MYB* gene alone or in combination with *MaMYC* (*MaMYB* + *MaMYC*). Intriguingly, overexpression of *MaMYBPAs* or co-expression of different combinations of *MaMYBs* and *MaMYCs* differentially affected the expression of *MaMYBPRs* (Figure 7f), suggesting that these repressors prevent excessive proanthocyanidin accumulation by forming a negative feedback loop.

### MaMYBPR interacts with different regulators in MBW complexes

The above results supported a physical interaction between MaMYBPAs and bHLH proteins to form a MBW complex. Since all MaMYBPRs also carry a bHLH motif in their N termini, we tested for an interaction between MaMYBPRs and their putative bHLH partners by yeast two-hybrid (Y2H) and BiFC assays. In Y2H, MaMYBPR1 interacted with MaMYC7 and MaMYC9, while MaMYBPR2, MaMYBPR3, and MaMYBPR4 interacted with MaMYC2, MaMYC7, and MaMYC9 (Figure 8a). We detected a stronger interaction between MaMYBPR1 and MaMYC7 and between MaMYBPR3 and MaMYC2, as evidenced by the appearance of blue colonies on selective medium with X-Gal, in contrast to the other combination for which the colonies remained white (Figure 8a). We confirmed the Y2H results by *in planta* BiFC assays in *N. benthamiana* leaves (Figure 8b). We failed to detect an *in planta* interaction for the combinations MaMYBPR1 and MaMYC9, MaMYBPR2-4 and MaMYC2-MaMYC7, or MaMYBPR3 and MaMYC7-MaMYC9, although we observed an interaction in Y2H, which may reflect different environments between the yeast and plant nucleus. These results indicate that MaMYB repressors might inhibit the activity of the MBW complex by interacting with MabHLH cofactors. We also analyzed whether these interacting combinations are co-localized and found that both MaMYBPR-MaMYC co-localized in the nucleus (Fig. 8c).

**Figure 8.**
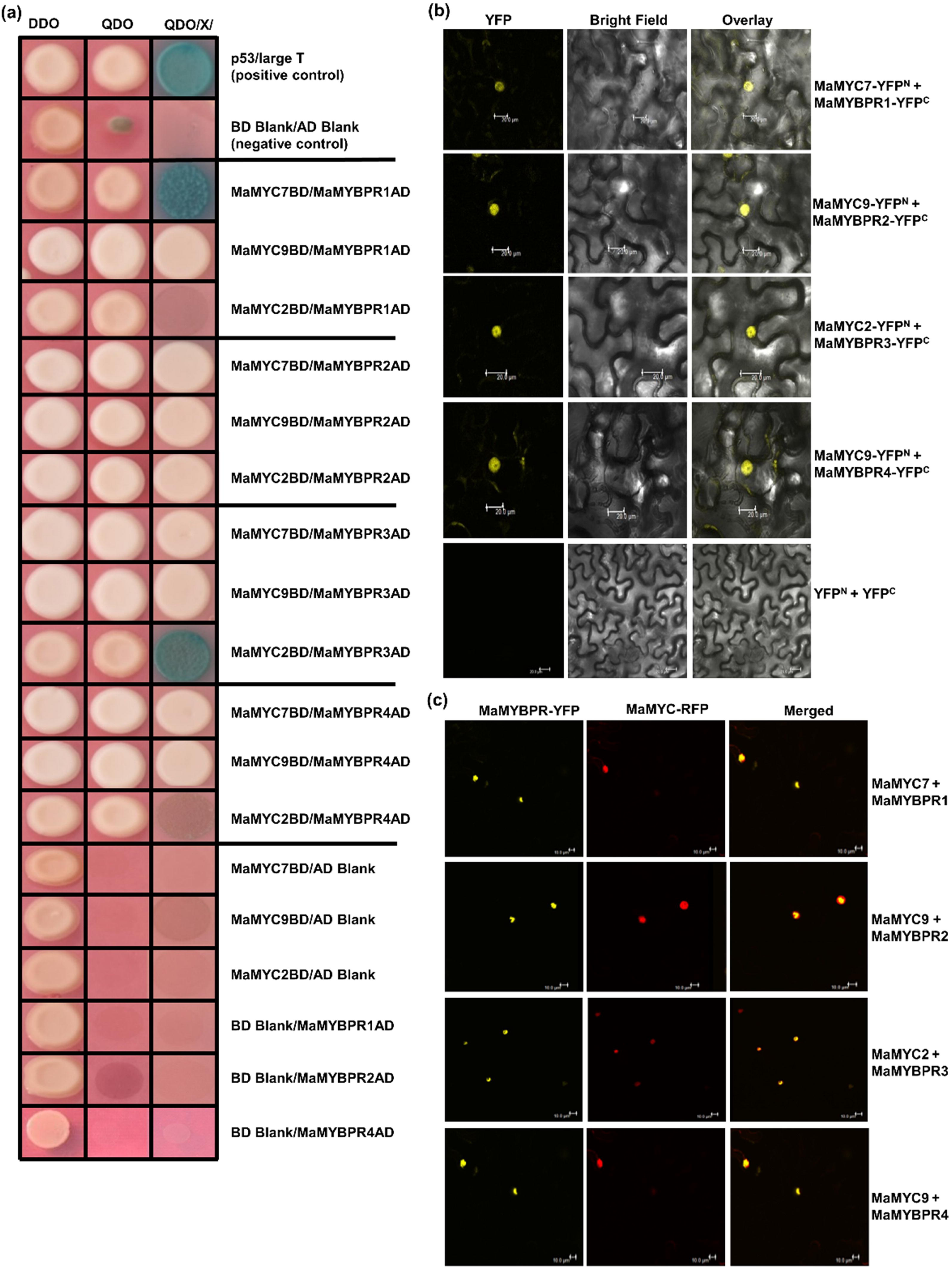
Interaction of R2R3 MYB repressors with MYC proteins of *M. acuminata*. (a) Yeast two-hybrid assays indicating the interaction of MaMYBL1-4 with MaMYC proteins. AD, GAL4 activation domain; BD, GAL4 DNA-binding domain; DDO, double synthetic defined (SD)–Leu–Trp medium; QDO/X/A, quadruple SD–Leu–Trp–Ade–His +X-α-Gal +AbA medium. (b) BiFC assays in transiently infiltrated *N. benthamiana* leaves indicating the *in planta* interaction of MaMYBL1-4 with MaMYC proteins. *YFP^N^ + YFP^C^* constructs were used as a negative control. Scale bars = 20 µm. (c) MaMYBL1-L4-RFP and MaMYB-YFP colocalize in the nucleus of transiently infiltrated *N. benthamiana* leaves, as analysed by confocal microscopy. Scale bars = 10 µm.

### MaMYBPR prevents proanthocyanidin accumulation in banana fruits by repressing proanthocyanidin biosynthesis genes

We ascertained the function of MaMYBPRs using transient expression assays in immature banana fruits. Accordingly, we overexpressed *MaMYBPR1–MaMYBPR4* individually or in combination with the corresponding *MaMYC* using the *ZmUBI1* promoter. Transformed fruits showed GUS activity, indicating successful transformation (Figure 9a). Spectrophotometric quantification of proanthocyanidins suggested that fruits transiently overexpressing *MaMYBPR1*, *MaMYBPR2*, *MaMYBPR3*, or *MaMYBPR4* individually or in combination with their *MaMYC* partners accumulate less proanthocyanidins compared to untransformed (control) fruit slices (Figure 9b). Relative transcript levels of the activator genes *MaMYBPA1* and *MaMYBPA2* and the biosynthesis genes *MaANS*, *MaLAR*, and *MaANR* were much lower in *MaMYBPR3*, *MaMYBPR4*, and *MaMYBPR1+MaMYC7* transformed fruits than in control fruit slices (Figure 9c and d). By contrast, fruits transiently overexpressing *MaMYBPR1* or *MaMYBPR2* alone or co-overexpressing *MaMYBPR2* and *MaMYC9*, or *MaMYBPR4* and *MaMYC9* showed increased expression levels for the same genes relative to control fruit slices (Figure 9c and d). We hypothesize that the activator genes *MaMYBPA1* and *MaMYBPA2* may be expressed to high levels in fruits overexpressing *MaMYBPR1*, *MaMYBPR2*, *MaMYBPR2*, and *MaMYC9*, or *MaMYBPR4* and *MaMYC9*, resulting in diminished overall repressor activity.

**Figure 9.**
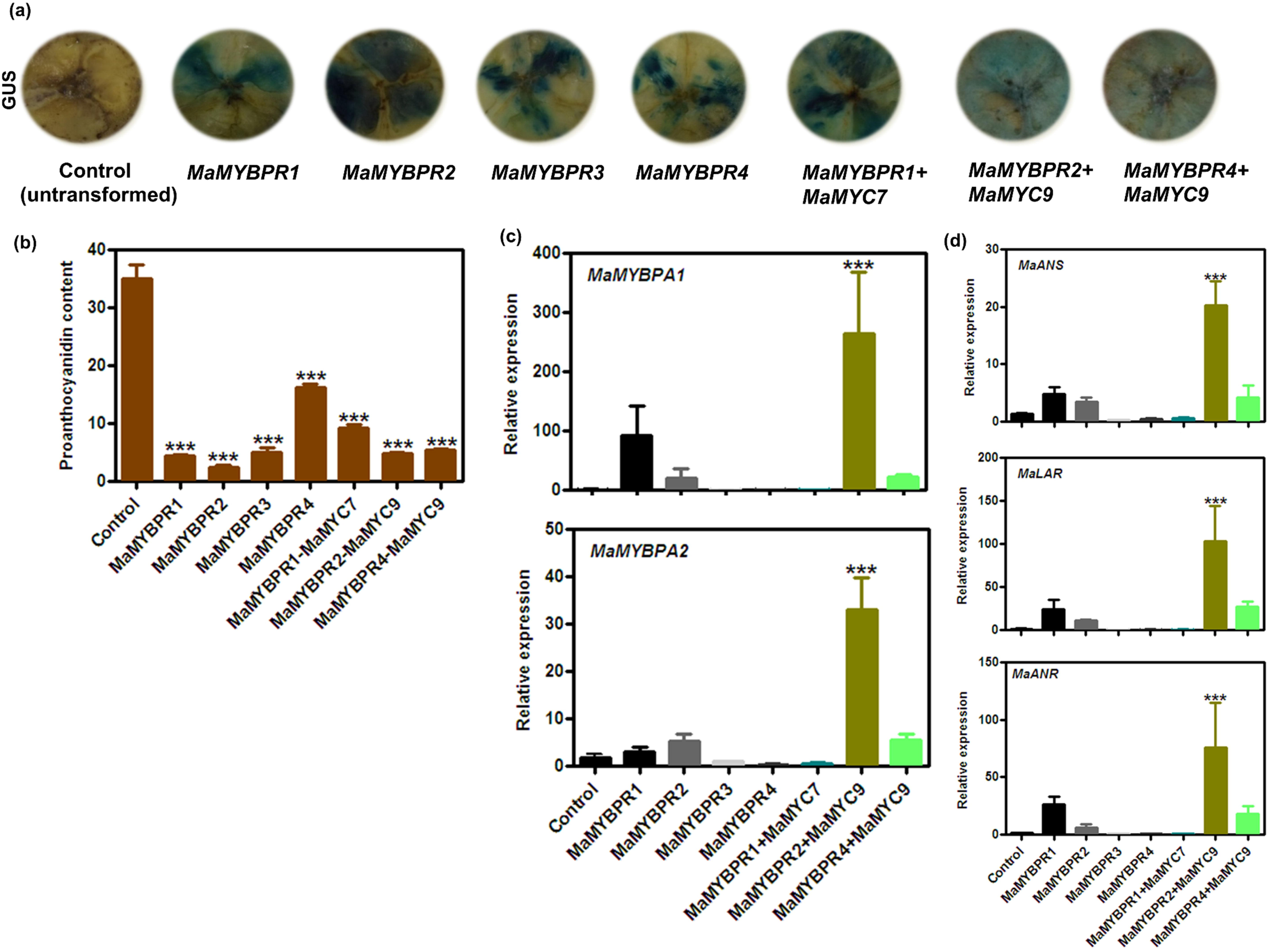
Functional analysis of MaMYB repressors in banana fruits. (a) Transiently transformed banana fruit slices with overexpression constructs of *MaMYBPR1-PR4* and *MaMYC*. Successful transformation is indicated by GUS reporter activity. (b) PA contents of transiently transformed fruit slices and control (empty vector). (c, d) Relative *MaMYBPA*s (c) and *MaANS*, *MaANR*, and *MaLAR* (d) transcript levels, as determined by RT-qPCR. Data are shown as means ±SD of three biological and three technical replicates (c, d). *, *p* ≤ 0.05; **, *p* ≤ 0.01, ***, *p* ≤ 0.001, as determined by one-way ANOVA.

### MaMYBPR physically interacts with MaMYBPA proteins

We tested the potential for interaction between MaMYBPRs and MaMYBPAs by BiFC assay in *N. benthamiana* leaves. We determined that MaMYBPA1 and MaMYBPA2 exhibit a TT8- independent interaction with MaMYBPR3 and MaMYBPR1, respectively, while neither MaMYBPA1 nor MaMYBPA2 interacted with MYBPR2 or MYBPR4 (Figure S6). These results indicate that PR-like MYB TFs physically interact with MaMYBPAs without requiring a bHLH partner for their incorporation into the MBW complex.

## Discussion

Despite our growing understanding of R2R3 MYB proteins as regulators of the proanthocyanidin biosynthesis, little is known about the fine-tuning regulatory loops that modulate proanthocyanidin accumulation in plants. The proanthocyanidin biosynthesis pathway responds to a variety of developmental and environmental cues, indicating the presence of a regulatory cascade involving both positive and negative regulators. In the present study, we identified and characterized two proanthocyanidin-specific R2R3-MYB activators and four R2R3-MYB repressors in banana. The MaMYBPAs is classified as a subgroup 5 R2R3-MYB TF family, grouped together all previously known positive regulators of proanthocyanidin biosynthesis (Verdier *et al*., 2012; Nesi *et al*., 2001; Schaart *et al*., 2013).

*MaMYBPA1* and *MaMYBPA2* expression in different organs of banana, particularly in fruit pulp, could be partially correlated with that of proanthocyanidin-specific structural biosynthesis genes expression, supporting the notion regarding their involvement in the transcriptional regulation of proanthocyanidin biosynthesis. We detected higher transcript levels for the regulators and biosynthesis genes in early stages of fruit pulp development, while proanthocyanidin-related metabolites were abundant in both leaves and peel tissues, indicating a lack of correlation between proanthocyanidin-related metabolites and transcript levels in leaf and peel tissues. In general, despite higher transcript abundance of biosynthesis enzyme encoding genes, lesser accumulation of metabolite hints at the possible proteolytic regulation of biosynthesis genes in these tissues. This can be exemplarily seen in *A. thaliana*, were the protein KFB^CHS^, a Kelch domain-containing F-box protein, physically interacts with the CHS protein, the key enzyme for the flavonoid biosynthesis, and specifically mediates its ubiquitination and degradation (Zhang *et al*., 2017). These findings also suggest that in addition to components of MBW complex, some additional upstream or parallel regulators might be working to regulate proanthocyanidin biosynthesis. For example apple protein MdWRKY41, a WRKY-type protein that suppresses the expression of *MdMYB12*, resulting in the repression of proanthocyanidin biosynthesis in apple (Mao *et al.,* 2021). In another example, *PpJMJ25*, a histone H3K9 demethylase gene in poplar epigenetically regulates the expression of an anthocyanin regulatory gene, resulting in a substantial drop in anthocyanin contents. PpJMJ25 demethylates H3K9me2 in the chromatin body of Pp*MYB182* to enhance its expression, leading to the downregulation of anthocyanin biosynthesis (Fan *et al.,* 2018). Since EAR motif by forming complex with chromatin-altering proteins regulates the expression of different genes at epigenetic level (Kagale and Rozwadowski, 2011), and in our analysis, the C2 motif, a type of EAR motif is present in all identified MaMYBPR proteins, such chromatin-altering proteins might form a complex with these repressors to modulate their repressor activity in banana. The grapevine gene *VvMYBPA1* transcripts are targeted by microRNA164c (miR164c) for mRNA cleavage, resulting in lower expression of *LAR* and *ANR*, and thus lower proanthocyanidin accumulation in grape berries (Vale *et al*., 2021). The varying accumulation of the transcripts of regulatory *R2R3-MYB* genes in banana tissues may be attributed to such regulators acting at different levels, which are yet to be identified. Therefore, in addition to transcriptional regulation, regulation at the post-transcriptional and at translational level would be interesting to study in future. Also, R2R3 MYB proteins act combinatory with other TF protein classes, which otherwise necessary for the activation of their target genes, making it difficult to correlate transcript levels of regulator genes with their targets. In the present study, we provide evidence that MaMYBPA1 and MaMYBPA2 interact with MaMYC and MaTTG1 proteins forming functional MBW complexes that were able to interact with the promoters of *MaANS, MaANR* and *MaLAR* and active the expression of these genes.

MYB- and bHLH-binding *cis*-motifs in the promoters of structural genes are required for MYB TF bindings. In persimmon fruit, DkMYB4 binds to the MYBCORE motif (CNGTTR) present in the promoters of the proanthocyanidin pathway genes *LAR* and *ANR* (Akagi *et al*., 2009). We also identified putative MYB binding sites (MBS), G-boxes (light responsiveness) known as bHLH binding sites in the *MaANS*, *MaLAR*, and *MaANR* promoters. In addition, we identified other *cis*-binding regulatory elements like ABRE sites (abscisic acid response element) and LTR motifs (low-temperature response) in these promoters, suggesting that different signals activate the expression of different structural genes in the flavonoid pathway. For example, FaRAV1, an ABA-responsive transcription factor, regulates anthocyanin biosynthesis by interacting with and increasing the expression of the CHS, F3H, DFR, and GT1 promoters (Zhang *et al*., 2020). In another example, light responsive transcription factors PIF3 and HY5 activates the expression of anthocyanin specific biosynthesis genes by binding to their light responsive units and positively control anthocyanin biosynthesis (Shin *et al*., 2007).

The heterologous overexpression of *MaMYBPA* genes partially restored proanthocyanidin accumulation in an Arabidopsis mutant lacking proanthocyanidins in its seeds, suggesting that MaMYBPA is a banana orthologue of Arabidopsis TT2. Overexpression of *MaMYBPA* genes individually or together with their interacting bHLH in banana fruits also activated proanthocyanidin biosynthesis, resulting in substantial proanthocyanidin accumulation, further validating MaMYBPA function. Moreover, the overexpression of R2R3 activator *MaMYBPA*s induced the expression of *R2R3 MYB* repressor genes, thus forming a negative feedback loop to modulate proanthocyanidin accumulation. Similarly, in peach, the expression of the repressor gene *PpMYB18* was induced by both the anthocyanin-related MYB activator PpMYB10.1 and the proanthocyanidin-related MYB activator PpMYBPA1 (Zhou *et al*., 2019). In another example, the proanthocyanidin-related activator MtMYB5 was shown to activate the transcription of the repressor gene *MtMYB2* in *M. truncatula* (Jun *et al*., 2015). It seems that this kind of regulation is a general principle for the fine-regulation of late flavonoid biosynthesis branches, which is found in dicots and also in monocots.

We have also identified and functionally characterized *MaMYBPR* repressor genes in banana. In our study, all four MYB repressors grouped together with other known MYB repressors from subgroup 4, which all comprise a R2R3 domain and a characteristic C1, C2, and TLLFR motif. The C2 motif, a type of EAR motif, is thought to work together with chromatin-remodelling complexes like DNA or histone methylases, demethylases, or acetyltransferases to modulate gene expression epigenetically (Kagale & Rozwadowski, 2011), suggesting potential epigenetic regulation of proanthocyanidin accumulation. Recently, MaMYBPR2 was identified as MaMYB4 which negatively regulate the anthocyanin biosynthesis in banana fruit and being a substrate of the RING-type E3 ligase MaBRG2/3 involved in fruit ripening (Deng *et al*., 2021; Yang et al. 2022).

We demonstrated here that the physical interaction between MaMYBPR, MaMYBPA, and MaMYC negatively modulates promoter activity in their target genes. MaMYBPR may interfere with the DNA binding ability of MaMYBPA, since MaMYBPA and MaMYBPR physically interact in a BiFC assay. Furthermore, overexpression of *MaMYBPR* alone or together with their interacting bHLH partners decreased the accumulation of proanthocyanidins in banana fruits, validating the function of MaMYBPR as a proanthocyanidin repressor. The accumulation of the repressor protein in banana unripe fruit appears to prevent overaccumulation of proanthocyanidins. However, the abundance of the repressor later diminishes with the onset of ripening since repressor seems to be regulated post-translationally by degradation via ubiquitin (Yang et al. 2022), leading to the accumulation of proanthocyanidins in ripe fruits.

Notably, *MaMYBPA* and downstream genes show modulation in their expression. In poplar, the repressive activity of PtMYB182 decreased when its bHLH-binding site was disrupted. In addition, poplar MYB182 was repressed by bHLH079 (GL3-type) or bHLH131 (TT8-like) cofactors to variable degrees (Yoshida *et al*., 2015). This observation is consistent with our expression data, in which the expression of biosynthetic genes and regulators is dependent on the bHLH protein and changes depending on the type of bHLH cofactor present. Our protein–protein interaction study demonstrated an interaction between the MaMYBPR repressors and MaMYC proteins to form another competitive MBW complex, which diminish the expression of proanthocyanidin biosynthesis genes. These findings support the relevance of the interaction with the bHLH cofactor for repressor activity, suggesting that MaMYBPRs may function in part by competing with activator MaMYBs for binding to bHLH, making the R3 domain of the MYB protein essential for repressor activity. Similar results were observed with peach PpMYB18 where amino acid substitutions in the R3 domain of PpMYB18 abolished its repressor activity (Zhou, 2019). In light of our findings, we propose a model that illustrates the interplay between activators and repressors and how activator induces the expression of repressor which later diminishes by degradation via ubiquitin to enhance proanthocyanidin biosynthesis in banana (Figure. 10). This study advanced our knowledge of the molecular basis of proanthocyanidin biosynthesis in monocots, in which proanthocyanidin regulation is less explored than in dicots and offers targets involved in proanthocyanidin biosynthesis for genome editing to further guide metabolic fluxes towards anthocyanins for banana biofortification. Since banana fruits are consumed raw, they would make an ideal vehicle for biofortification of thermolabile anthocyanins.

**Figure 10.**
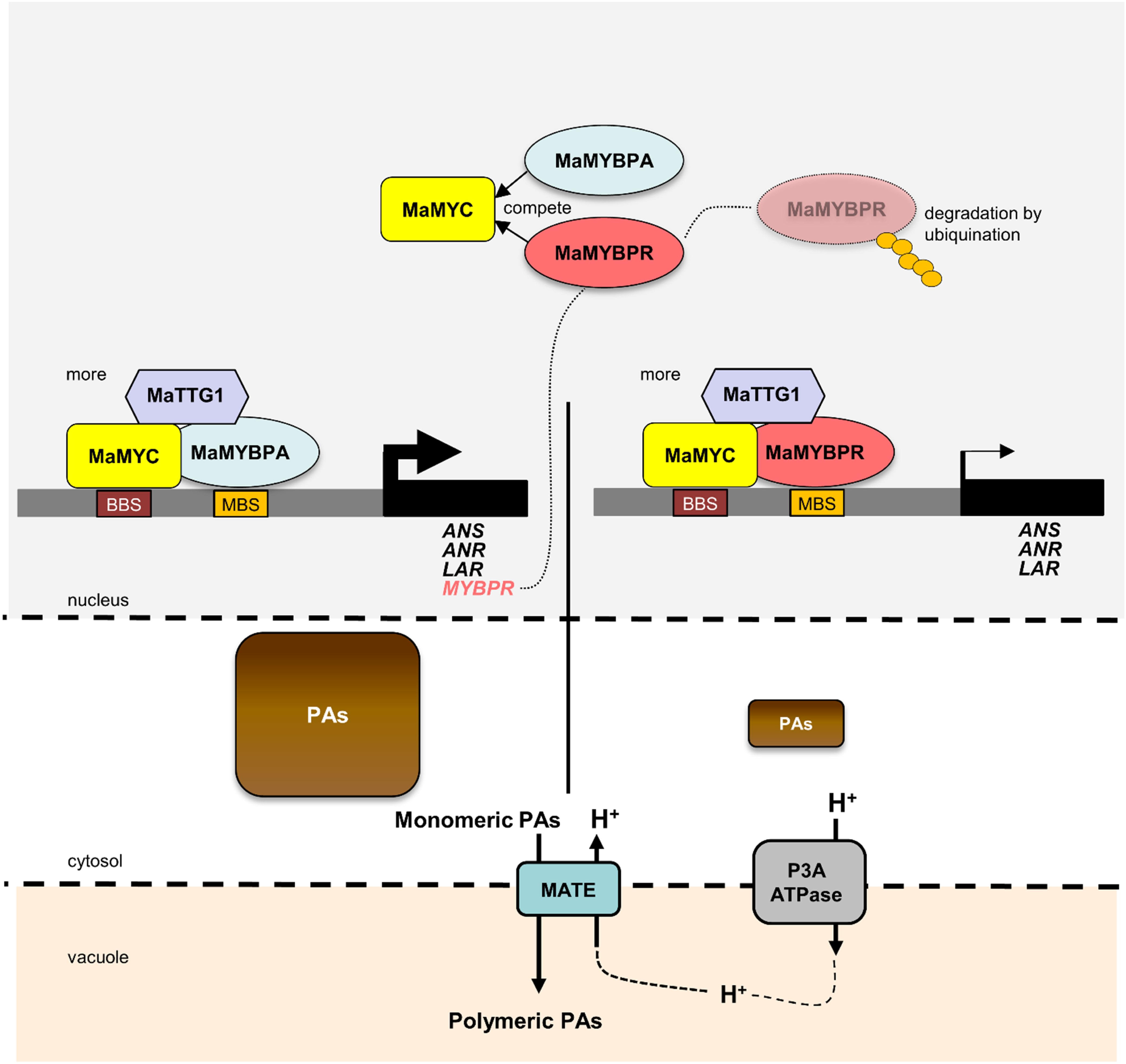
Working model of the interplay between MYB activators and repressors to balance the proanthocyanidin biosynthesis in banana fruit. MaMYBPA together with MaMYC and MaTTG1 directly bind to the MBS and BBS sites of target gene promoters and activate transcription (marked by solid black arrow). MaMYBPR together with MaMYC inhibits the formation of the MBW complex, which leads to the repression (marked by fine black arrow) of target gene expression. We also hypothesize the role of MATE transporters in proanthocyanidin transport from the cytosol to the vacuole. MBS, MYB binding site; BBS, bHLH binding site.

## Supporting information

Supplementary Table S1 and S2. Supplementary Fif. S1 to S6

## Acknowledgements

This work was supported by the core grant of National Institute of Plant Genome Research and Department of Science and Technology-SERB for Startup research grant to AP. RR and JN acknowledges Council of Scientific and Industrial Research, Government of India, for Senior Research Fellowships. The authors are thankful to DBT-eLibrary Consortium (DeLCON) for providing access to e-resources. We acknowledge the Metabolome facility (BT/ INF/22/SP28268/2018) at NIPGR for phytochemical analysis.

## Author’s contribution

AP conceived the idea and designed the research. RR and JN conducted experiments. RR and AP interpreted the data. RR, RS and AP wrote the manuscript. All authors read and approved the final manuscript.

## Data availability

All data supporting the findings of this study are available within the paper and within the Supplementary data published online.

## Declarations

The authors declare no conflict of interest.

## Supporting Information

**Figure S1** R2R3-MYB repressors from M. acuminata (MaMYBPR1 to MaMYBPR4).

**Figure S2** Representative UHPLC and MS-MS chromatograms.

**Figure S3** Identification and expression analysis of candidate bHLH proteins involved in proanthocyanidin biosynthesis from *M. acuminata*.

**Figure S4** Co-localization of MaMYBPA and MaMYC.

**Figure. S5** Phylogeny of TTG1-like proteins.

**Figure. S6** Interaction of MaMYBPA proteins with MaMYB repressors.

**Table S1** Accession number of protein sequences used for multiple sequence alignment and phylogenetic analysis.

**Table S2** List of primers used in the present study.

