## Supplementary Table S1 and S2. Supplementary Fif. S1 to S6 for "Interplay between R2R3 MYB-type activators and repressors regulates proanthocyanidin biosynthesis in banana (*Musa acuminata*)"

Supporting information = 06 (Figures); 02 (Table)

Colored Supporting information = Figure S1-S6

**ORCID**

Ruchika Rajput (0000-0003-4103-1973), Jogindra Naik (0000-0002-3435-6502), Ralf Stracke, (0000-0002-9261-2279), Ashutosh Pandey, (0000-0002-5018-9628)

**Supporting Method:**

**Ultra-High-Performance Liquid Chromatography (UHPLC)**

Separation for qualitative and quantitative analysis of monomeric and oligomeric PAs was performed on a 1290 Infinity II series HPLC system (Agilent Technologies) equipped with a 1290 Infinity II series pump, autosampler, column compartment, and thermostat using a Zorbax Eclipse Plus $C_{18}$ column (2.1 x 100 mm, 1.8 µm) maintained at 30°C. The mobile phase consisted of an aqueous solution of 0.1% (v/v) LC-MS-grade formic acid (solution A) and 0.1% formic acid in LC-MS-grade acetonitrile (solution B). The gradient for solution B was programmed as follows: 0 to 2 min, 5% (v/v); 2 to 10 min, 5%–15%; 10 to 32 min, 15%-30%; 32 to 40 min, 30%-80%; 40 to 42 min, 80%–90%; and 42–47 min, 90%-0%. Other chromatographic parameters included a constant flow of 270 µL/min (injection volume, 3 µL), and run time of 47 min including equilibration.

**Liquid chromatography–mass spectrometry (LC-MS)**

Analysis of PAs was performed using an UPLC system (Exion LC Sciex) coupled to a triple quadrupole system (QTRAP6500+, ABSciex) using electrospray ionization (ESI). The voltage was set to 5,500 V for positive ionization. The following parameters were applied: gas 1 and gas 2 (70 psi), curtain gas (40 psi), collision-assisted dissociation (medium), and temperature of the source (650°C). The mass spectrometer was used in multiple reaction monitoring mode (MRM) for qualitative and quantitative analysis. Analytical standards were purchased from Merck (Germany). Identification and quantitative analysis were carried out using Analyst software version 1.5.2 (ABSciex).

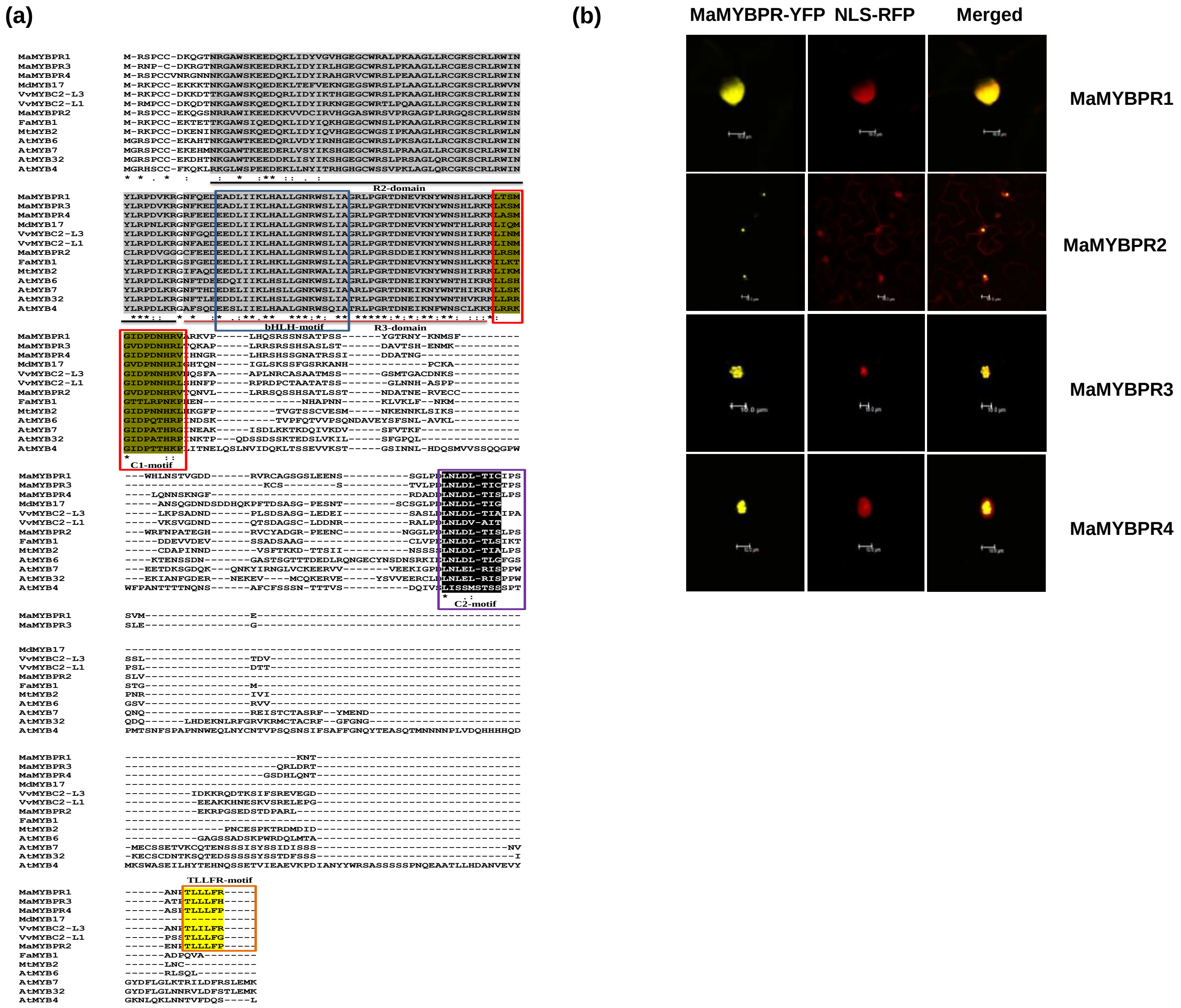


**Figure S1** R2R3-MYB repressors from *M. acuminata* (MaMYBPR1 to MaMYBPR4).: Multiple sequence alignment of R2R3-MYB-type repressors. (a)The R2 and R3 repeats of the MYB domain are the bHLH interaction motif highlighted in grey colour, C1 and C2 motif are given in green and black colour, respectively. The TLLFR motif is indicated in yellow according to previously characterized R2R3-MYB repressors. (b). Subcellular localization of MaMYBPR1 to PR4-YFP fusion proteins in Agrobacteria-infiltrated tobacco leaves, analysed by confocal microscopy. NLS-RFP was used as nuclear marker. Scale bar = 10 µm.

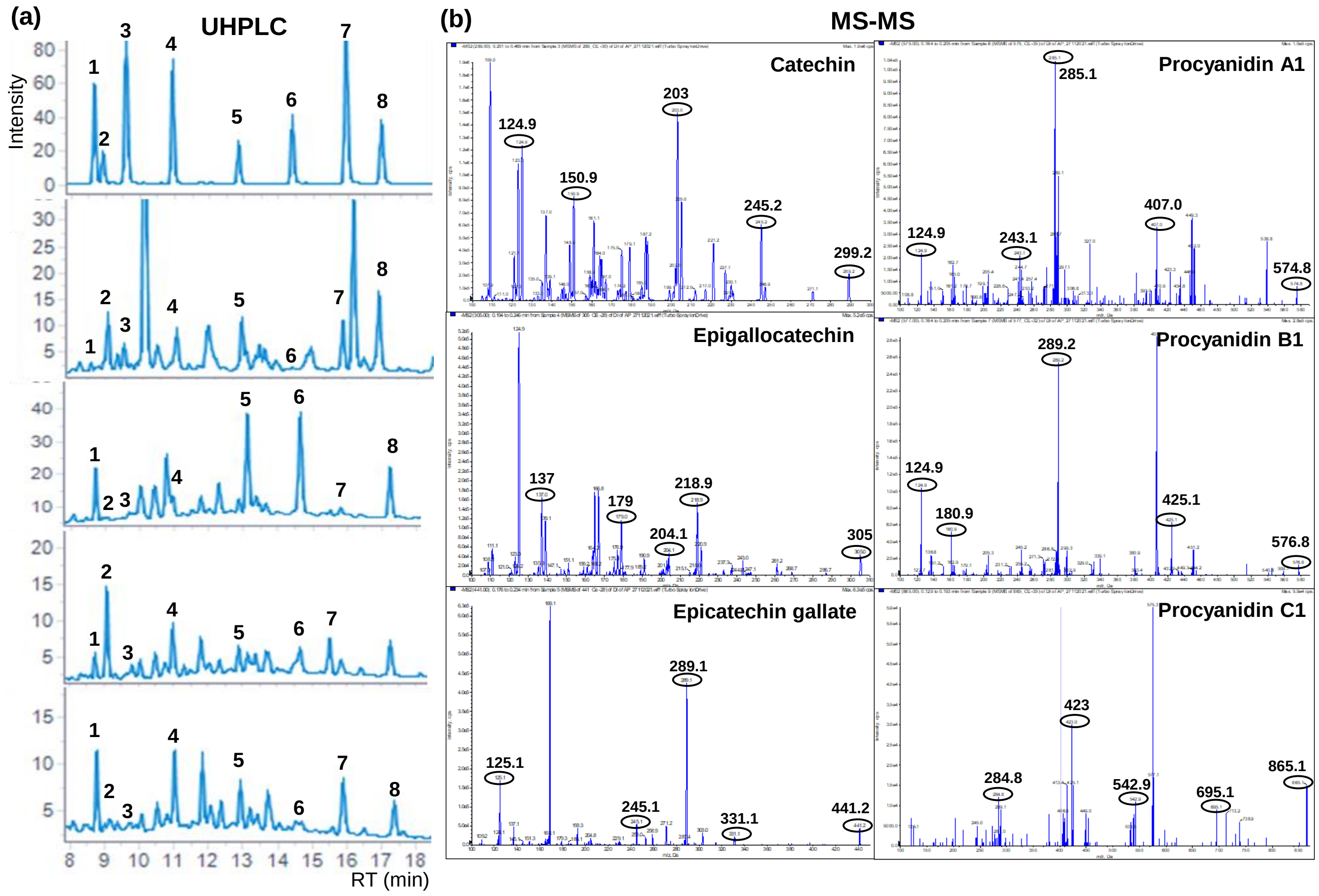


**Figure S2** Representative UHPLC and MS-MS chromatograms. Representative UHPLC chromatograms of standard PAs, and methanolic extracts of different organs of *M. acuminata* are shown in the figure. Different peaks in the chromatogram are of Procyanidin B1 (1); Epigallocatechin (2); Catechin (3); Procyanidin B2 (4); Procyanidin C1 (5); Procyanidin A1 (6); Epicatechin gallate (7); Procyanidin A2 (8) (a). MS-MS spectrum of representative monomeric (Catechin; Epigallocatechin and Epicatechin gallate) and oligomeric (Procyanidin A1; Procyanidin B1 and Procyanidin C1) PAs in negative mode (b).

(a)

(b)

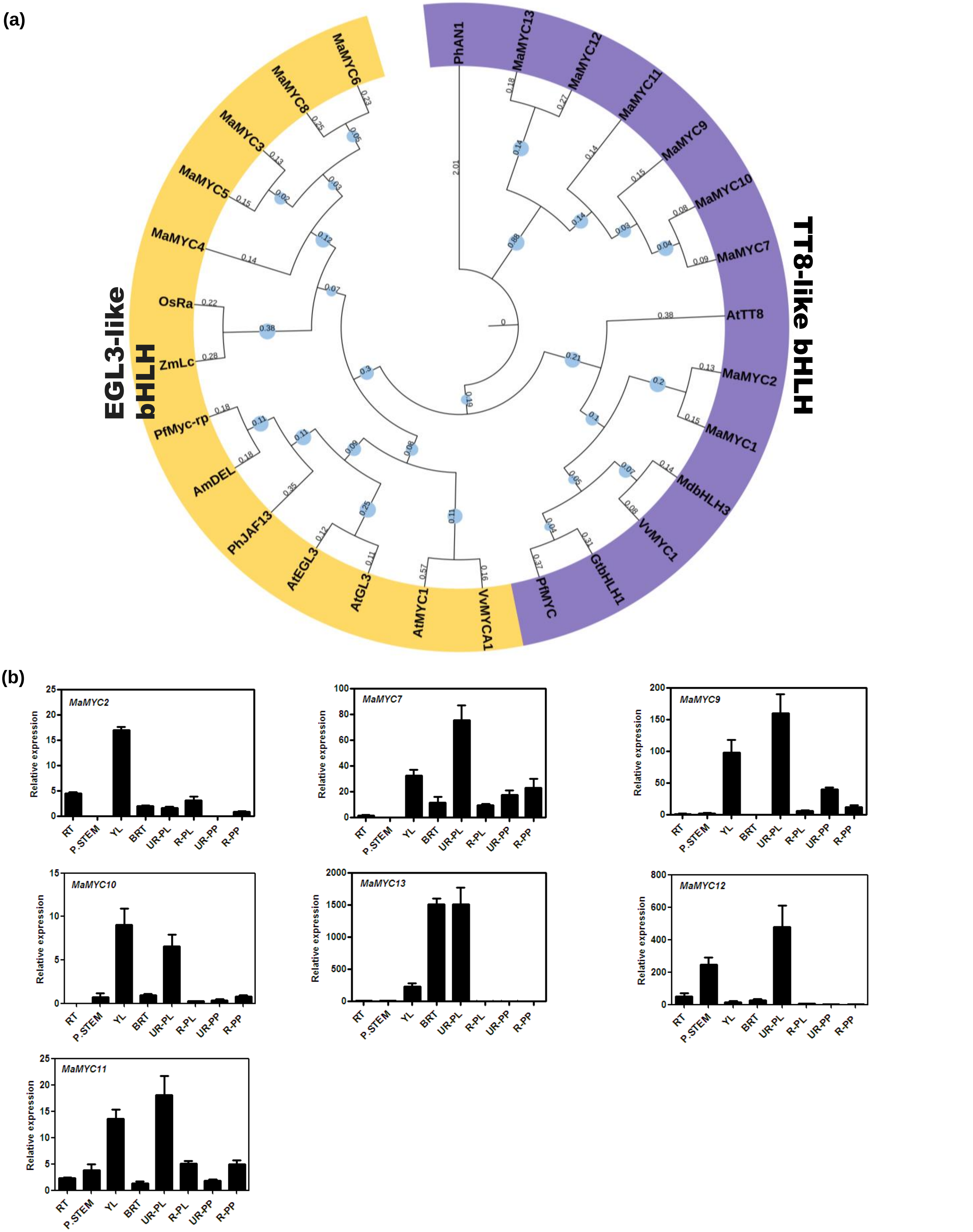


**Figure S3** Identification and expression analysis of candidate bHLH proteins involved in proanthocyanidin biosynthesis from *M. acuminata.* Cladogram of TT8- and EGL3-like bHLH proteins from *M. acuminata* and landmark bHLH-type PA regulators from other plant species. *M. acuminata* bHLH proteins were named as MaMYC1-13 ordered by their position on the pseudochromosomes. Bootstrap values are shown with light blue circles at the nodes, indicating higher values in larger circles. Numbers on each branch indicate the branch length (a). RT-qPCR expression analysis of TT8-like *MaMYC* genes were carried out in YL (young leaves), BRT (bract), P.STEM (pseudostem), RT (root), UR-PL (unripe peel), UR-PP (unripe pulp), R-PL (ripe peel) R-PP (ripe pulp).. The graphs show values ±SD of three technical replicates. *MaActin* was used as reference control (b).

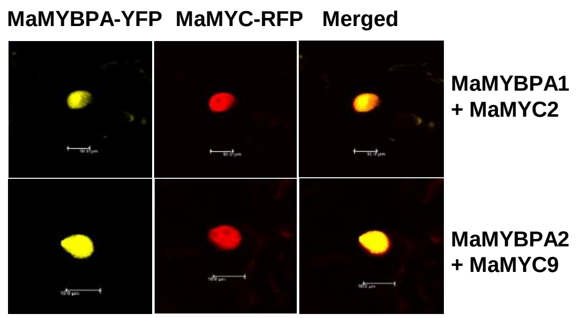


**Figure S4** Co-localization of MaMYBPA and MaMYC. MaMYBPA-YFP and MaMYC-RFP are co-localized in the nucleus of Agrobacteria-infiltrated tobacco leaves analysed by confocal microscopy. Scale bar = 10 µm.

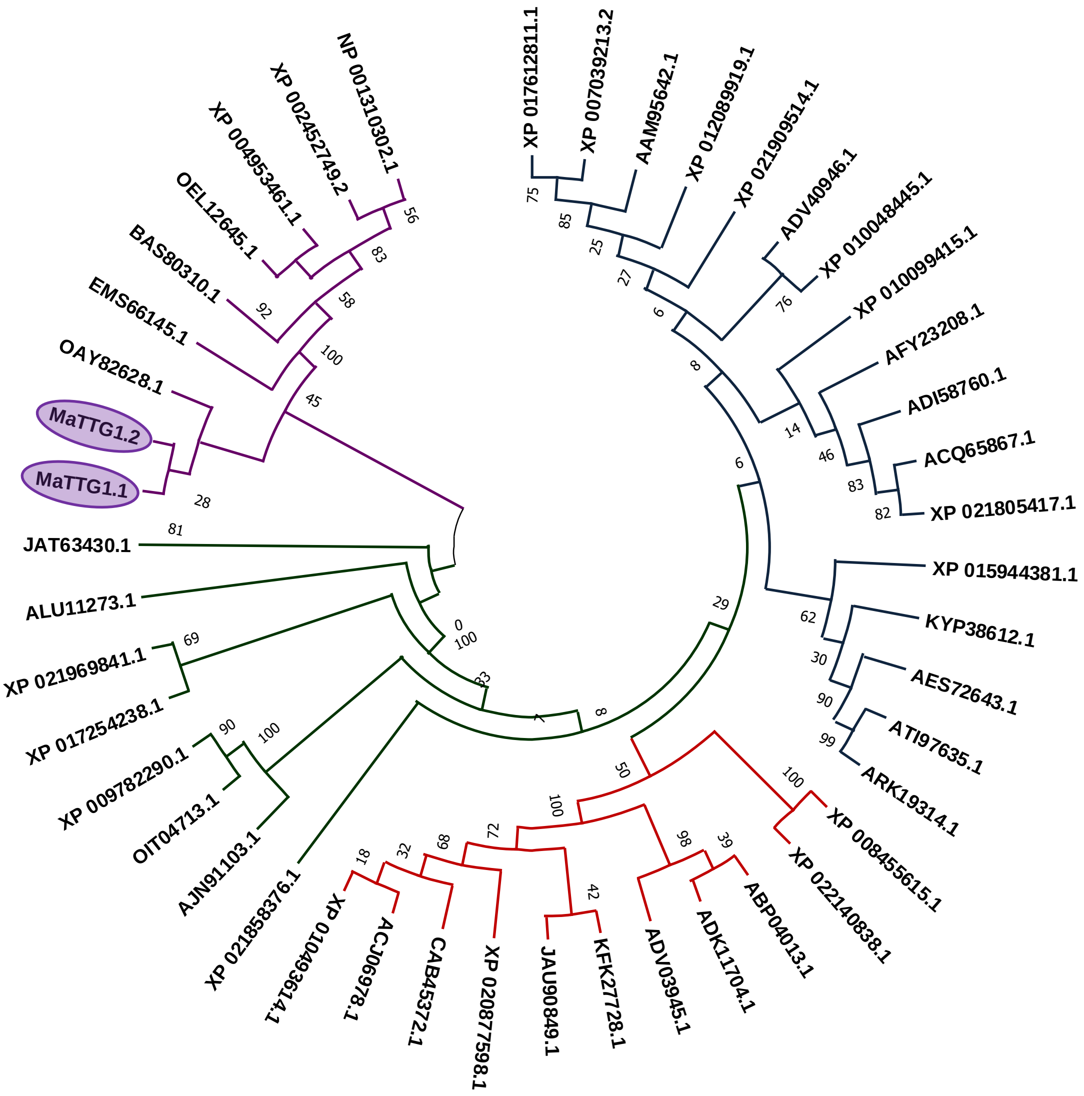


**Figure S5** Phylogeny of TTG1-like proteins. *Musa acumintae* TTG1-like proteins (MaTTG1.1:Ma04_35650, MaTTG1.2 Ma05_09990) were employed for the construction of phylogeny using maximum likelihood (ML) method at 1000 bootstrap replication along with TTG1 proteins from *Ananas comosus* (OAY82628.1), *Anthurium amnicola* (JAT63430.1), *Arabidopsis lyrata* (XP_020877598.1), *A. thaliana* (CAB45372.1), *Arabis alpina* (KFK27728.1), *Arachis duranensis* (XP_015944381.1), *Brassica napus* (ABP04013.1), *Brassica oleracea* (ADV03945.1), *Brassica rapa* (ADK11704.1), *Cajanus cajan* (KYP38612.1), *Camelina sativa* (XP_010493614.1), *Carica papaya* (XP_021909514.1), *Cucumis melo* (XP_008455615.1), *Daucus carota* (XP_017254238.1), *Dichanthelium oligosanthes* (OEL12645.1), *Eucalyptus grandis* (XP_010048445.1), *Gossypium arboreum* (XP_017612811.1), *G. hirsutum* (AAM95642.1), *Helianthus annuus* (XP_021969841.1), *Jatropha curcas* (XP_012089919.1), *Lotus corniculatus* (ARK19314.1), *L. sessilifolius* (ATI97635.1), *Malus domestica* (ADI58760.1), *Medicago truncatula* (AES72643.1), *Momordica charantia* (XP_022140838.1), *Morus notabilis* (XP_010099415.1), *Nelumbo lutea* (ALU11273.1), *Nicotiana attenuata* (OIT04713.1), *N. sylvestris* (XP_009782290.1), *N. tabacum* (ACJ06978.1), *Noccaea caerulescens* (JAU90849.1), *Oryza sativa* (BAS80310.1), *Prunus avium* (XP_021805417), *P. persica* (ACQ65867.1), *Punica granatum* (ADV40946.1), *Rosa rugosa* (AFY23208.1), *Setaria italica* (XP_004953461.1), *Sorghum bicolor* (XP_002452749.2), *Solanum melongena* (AJN91103.1), *Spinacia oleracea* (XP_021858376.1), *Triticum urartu* (EMS66145.1), *Zea mays* (NP_001310302.1). *Musa acumintae* TTG1-like proteins Ma04_35650 and Ma05_09990 (blue highlighted) were closely clustered with TTG1 proteins from plants like *Ananas comosu, Anthurium amnicola, Nelumbo lutea, Triticum urartu.*

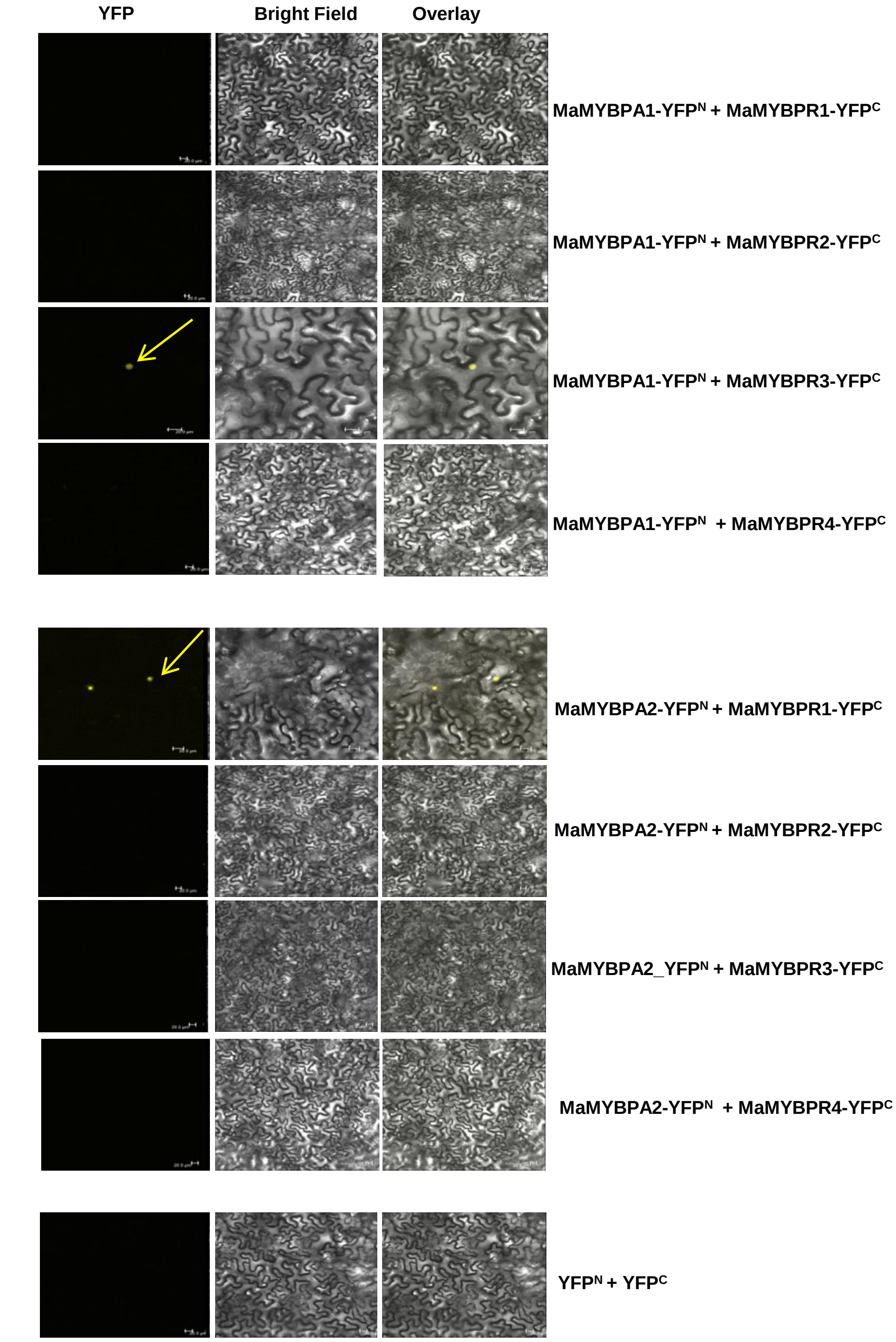


**Figure S6** Interaction of MaMYBPA proteins with MaMYB repressors. BiFC assays in Agrobacteria-infiltrated *N. benthamiana* leaves indicate *in planta* interaction of MaMYBPA1 with MaMYBPR3, and interaction of MaMYBPA2 with MaMYBPR1. $YFP^N$-$YFP^C$ was used as negative control. Scale bar =20 µm.

**Table S1: Accession number of protein sequences used for multiple sequence alignment and phylogenetic analysis**

The sequences of the R2R3-MYB domains were extracted from the full-length MaMYB protein sequences. The multiple sequence alignment of MaMYBPAs and MaMYBPRs was carried out with the functionally characterized R2R3-MYB-type PA regulator proteins

| Gene Name | Accession number |
|---|---|
| TT2 | NM122946 |
| MdMYB9 | DQ267900 |
| MdMYB11 | DQ074463 |
| MdMYB12 | XP008337875.1 |
| VvMYBPA2 | ACK56131.1 |
| FaMYB9 | JQ989281 |
| AtMYB7 | NP_179263 |
| AtMYB4 | At4g38620 |
| FaMYB1 | AF401220 |
| MdMYB17 | HM122618 |
| VvMYBC2-L3 | KM046932 |
| VvMYBC2-L1 | JX050227 |
| MtMYB2 | XM_003616340 |

**Table S2: List of primers used in the present study**

| Primer Name | Sequence (5’ to 3’) | Purpose |
|---|---|---|
| MaMYBPA1_For | 5’CGATGACTCGCTATGGTTT3’ | RT-qPCR forward primer |
| MaMYBPA1_Rev | 5’TAC CGGAAGTGGGATCTC3’ | RT-qPCR reverse primer |
| MaMYBPA2_For | 5’ACGACTTGATGTCCCTCG3’ | RT-qPCR forward primer |
| MaMYBPA2_Rev | 5’ACGCTGCAAACTGGAGAT3’ | RT-qPCR reverse primer |
| MaMYC2_For | 5’GGAGCTGAAGAAGGCGATTCA3' | RT-qPCR forward primer |
| MaMYC2_Rev | 5'AGTATTAGCGGGCCATTGCTG3' | RT-qPCR reverse primer |
| MaMYC7 _For | 5'CTCCGTGCTGCTCTCTACTC3' | RT-qPCR forward primer |
| MaMYC7 _Rev | 5'CCGCCATTAAAGGCCAAGAC3' | RT-qPCR reverse primer |

| | | |
|---|---|---|
| MaMYC9_For | 5'TCTCTACGCTAGATTGGCGG3' | RT-qPCR forward primer |
| MaMYC9_Rev | 5'CTCCATGGCGGCAGTAAGA3' | RT-qPCR reverse primer |
| MaMYC10_For | 5' GCGGAGGCACCAATTAGTAGT3' | RT-qPCR forward primer |
| MaMYC10_Rev | 5'AAAATTTCTCCCGCCGCCAT3' | RT-qPCR reverse primer |
| MaMYC11_For | 5'AACGCTGCTCTCTATTCCCG3' | RT-qPCR forward primer |
| MaMYC11_Rev | 5'AAGCGCAATACACAATACAGCA3' | RT-qPCR reverse primer |
| MaMYC12_For | 5'CCGAGTCGCATAGCCTTTTTC3' | RT-qPCR forward primer |
| MaMYC12_Rev | 5'TTGTTGTACACCCACCACCTT3' | RT-qPCR reverse primer |
| MaMYC13_For | 5'CTGATCATCCTCCTCACGCC3' | RT-qPCR forward primer |
| MaMYC13_Rev | 5'GGCAGTGGCTGCTCTAGATT3' | RT-qPCR reverse primer |
| AtLDOX_For | 5’GACGATGAAAAGATCCGTGAG3’ | RT-qPCR forward primer |
| AtLDOX_Rev | 5’CTCTTCTCCGGCTTTCTTG3’ | RT-qPCR reverse primer |
| AtBAN_For | 5’TTCATCACCGGGAAAGAAATG3’ | RT-qPCR forward primer |
| AtBAN_Rev | 5’AACAAATGGGCACGAGCTAAA3’ | RT-qPCR reverse primer |
| MaANS_For | 5'GAGGGGAAGCTGGACAGGGAAC3' | RT-qPCR forward primer |
| MaANS_Rev | 5'GTTGTGGAGGATGAAGGAGAGC3' | RT-qPCR reverse primer |
| MaLAR_For | 5'GCATTCTGGATCAGCTCTGCTTGC3' | RT-qPCR forward primer |
| MaLAR_Rev | 5'CTCGCCGAACCCTTCTCTTTCTG3' | RT-qPCR reverse primer |
| MaANR_For | 5’CCGGTTCCATCTCGTTCACCCAC3’ | RT-qPCR forward primer |
| MaANR_Rev | 5’CCGAGGGTATCGTTCTGAGAGGA3’ | RT-qPCR reverse primer |
| MaActin_For | 5'ATGACATGGAGAAGATCTGGCATCA3' | RT-qPCR forward primer |
| MaActin_Rev | 5'AGCCTGGATGGCAACATACATAGC3' | RT-qPCR reverse primer |
| MaMYBPR1_For | 5'GAGGACCAGAAGCTCATCGA3' | RT-qPCR forward primer |
| MaMYBPR1_Rev | 5'CAACTCTTACCGCACCGAAG3' | RT-qPCR reverse primer |
| MaMYBPR2_For | 5'CCCGATCTGAACCTTGACCT3' | RT-qPCR forward primer |
| MaMYBPR2_Rev | 5'ATGGAAAGAGGAGGAGCGTT3' | RT-qPCR reverse primer |
| MaMYBPR3_For | 5'TCCACTGTGTTGCCTGATCT3' | RT-qPCR forward primer |
| MaMYBPR3_Rev | 5'AGTCCTGTCAAGCCTCTGTC3' | RT-qPCR reverse primer |
| MaMYBPR4_For | 5'CGAGGAGGATGAAGAGGACC3' | RT-qPCR forward primer |
| MaMYBPR4_Rev | 5'TTGCCAGCTTCTTCCTCAGA3' | RT-qPCR reverse primer |
| MaMYBPA1_For | 5’GGGGACAAGTTTGTACAAAAAAGCAGGCTccATGGGGAGAAGGCCTTGCTGT3’ | attB1 forward primer for full length cloning of MaMYBPA1_ cDNA into |

| | | |
|---|---|---|
| | | entry clone |
| MaMYBPA1_Rev | 5’GGGGACCACTTTGTACAAGAAAGCTGGGTaTTTGTCTTGCGCTTCTTCTTCA3’ | attB2 reverse primer for full length cloning of MaMYBPA1_ cDNA into entry clone |
| MaMYBPA2_For | 5’GGGGACAAGTTTGTACAAAAAAGCAGGCTccATGGGAAGGAAGCCTTGTTGTT 3’ | attB1 forward primer for full length cloning of MaMYBPA2_ cDNA into entry clone |
| MaMYBPA2_Rev | 5’GGGGACCACTTTGTACAAGAAAGCTGGGTaGCTCCAATACCCCTGTTCGTCA3’ | attB2 reverse primer for full length cloning of MaMYBPA2_ cDNA into entry clone |
| MaMYC2_For | 5’GGGGACAAGTTTGTACAAAAAAGCAGGCTccATGGTCGCCCCACCGAGCACCGGTC 3' | attB1 forward primer for full length cloning of MaMYC2_ cDNA into entry clone |
| MaMYC2_Rev | 5'GGGGACCACTTTGTACAAGAAAGCTGGGTaATGGTCAGAGATTATTTGGTGAATC 3' | attB2 reverse primer for full length cloning of MaMYC2_ cDNA into entry clone |
| MaMYC3_For | 5’GGGGACAAGTTTGTACAAAAAAGCAGGCTccATGGAAGCAGATCTCGAAATCCATG 3' | attB1 forward primer for full length cloning of MaMYC3_ cDNA into entry clone |
| MaMYC3_Rev | 5'GGGGACCACTTTGTACAAGAAAGCTGGGTaCAGGCACTTGCTTATCACTCTCTG 3' | attB2 reverse primer for full length cloning of MaMYC3_ cDNA into entry clone |
| MaMYC7_For | 5’GGGGACAAGTTTGTACAAAAAAGCAGGCTccATGCGCAGGAGTCGGTGCCCGGTGTT 3' | attB1 forward primer for full length cloning of MaMYC7_ cDNA into entry clone |
| MaMYC7_Rev | 5'GGGGACCACTTTGTACAAGAAAGCTGGGTaCCTGCTAATTGGTGCCTCGGCAGCCA3' | attB2 reverse primer for full length cloning of MaMYC7_ cDNA into entry clone |
| MaMYC9_For | 5’GGGGACAAGTTTGTACAAAAAAGCAGGCTccATGAACCTGTGGGCCGACGACAA3' | attB1 forward primer for full length cloning of MaMYC9_ cDNA into entry clone |
| MaMYC9_Rev | 5'GGGGACCACTTTGTACAAGAAAGCTGGGTaCCAACTAGATAAGGCCTCAGCC3' | attB2 reverse primer for full length cloning of MaMYC9_ cDNA into entry clone |
| MaMYC13_For | 5’GGGGACAAGTTTGTACAAAAAAGCAGGCTccATGCAACGGTCGGATCACCG3' | attB1 forward primer for full length cloning of MaMYC13_ cDNA into entry clone |
| MaMYC13_Rev | 5'GGGGACCACTTTGTACAAGAAAGCTGGGTaCATGGCGTGAGGAGGATGATCAGCT3' | attB2 reverse primer for full length cloning of MaMYC13_ cDNA into entry clone |
| MaMYBPR1_For | 5’GGGGACAAGTTTGTACAAAAAAGCAGGCTccATGAGGAGCCCTTGTTGTGATA | attB1 forward primer for full length cloning of |

| | | |
|---|---|---|
| | 3’ | MaMYBPR1_ cDNA into entry clone |
| MaMYBPR1_Rev | 5'GGGGACCACTTTGTACAAGAAAGCTGGGTaTTAGCGAAAGAGAAGGAGCGTC3’ | attB2 reverse primer for full length cloning of MaMYBPR1_ cDNA into entry clone |
| MaMYBPR2_For | 5’GGGGACAAGTTTGTACAAAAAAGCAGGCTccATGAGGAGTCCTTGTTGTGAGA 3’ | attB1 forward primer for full length cloning of MaMYBPR2_ cDNA into entry clone |
| MaMYBPR2_Rev | 5'GGGGACCACTTTGTACAAGAAAGCTGGGTaTTATGGAAAGAGGAGGAGCGTT3’ | attB2 reverse primer for full length cloning of MaMYBPR2_ cDNA into entry clone |
| MaMYBPR3_For | 5’GGGGACAAGTTTGTACAAAAAAGCAGGCTccATG AGG AAC CCT TGC GAC AAGC3’ | attB1 forward primer for full length cloning of MaMYBPR3_ cDNA into entry clone |
| MaMYBPR3_Rev | 5'GGGGACCACTTTGTACAAGAAAGCTGGGTa CAA TGA AAG AGA AGA AGC GTC3’ | attB2 reverse primer for full length cloning of MaMYBPR3_ cDNA into entry clone |
| MaMYBPR4_For | 5’GGGGACAAGTTTGTACAAAAAAGCAGGCTccATGAGGAGCCCTTGTTGCGTTA 3’ | attB1 forward primer for full length cloning of MaMYBPR4_ cDNA into entry clone |
| MaMYBPR4_Rev | 5'GGGGACCACTTTGTACAAGAAAGCTGGGTaTCATGGAAAGAGGAGGAGCGTA3’ | attB2 reverse primer for full length cloning of MaMYBPR4_ cDNA into entry clone |
| proMaANR_For | 5’GGGGACAAGTTTGTACAAAAAAGCAGGCTccGGCCCTCTGATCACAGTTACTTTC3’ | attB1 forward primer for cloning of MaANR promoter into entry clone for co-transfection and dual luciferase assay |
| proMaANR_Rev | 5’GGGGACCACTTTGTACAAGAAAGCTGGGTaCCACAGTAAAGCTTAACCACCGTT3’ | attB2 reverse primer for cloning of MaANR promoter into entry clone for co-transfection and dual luciferase assay |
| proMaLAR_For | 5’GGGGACAAGTTTGTACAAAAAAGCAGGCTccCACTTAGAACTTGAGGAGGATTCT 3’ | attB1 forward primer for cloning of MaLAR promoter into entry clone for co-transfection and dual luciferase assay |
| proMaLAR_Rev | 5’GGGGACCACTTTGTACAAGAAAGCTG | attB2 reverse primer for cloning of MaLAR promoter |

| | | |
|---|---|---|
| | GGTaGGCGGGGAGAATATGCTGAAA 3’ | into entry clone for co-transfection and dual luciferase assay |
| proMaANS_For | 5’GGGGACAAGTTTGTACAAAAAAGCAGGCTccTTTATACTCTTAGCACTACGATTTTGGG3’ | attB1 forward primer for cloning of MaANS promoter into entry clone for co-transfection and dual luciferase assay |
| proMaANS_Rev | 5’GGGGACCACTTTGTACAAGAAAGCTGGGTaTAGGAGAAGCTTCCGCTACTGGCAGCAA3’ | attB2 reverse primer for cloning of MaANS promoter into entry clone for co-transfection and dual luciferase assay |
| proMaANR_For | 5’GGGGACAAGTTTGTACAAAAAAGCAGGCTccCCATTTGCATGTAGATGATGTAGTTG3’ | attB1 forward primer for cloning of MaANR promoter into entry clone for Y1H assay |
| proMaANR_Rev | 5’GGGGACCACTTTGTACAAGAAAGCTGGGTaTCTATGATATGCAATACAAGATGGAC3’ | attB2 reverse primer for cloning of MaANR promoter into entry clone for Y1H assay |
| proMaLAR_For | 5’GGGGACAAGTTTGTACAAAAAAGCAGGCTccCTTTTGGGATACTTGGGGAACTGTG3’ | attB1 forward primer for cloning of MaLAR promoter into entry clone for Y1H assay |
| proMaLAR_Rev | 5’GGGGACCACTTTGTACAAGAAAGCTGGGTaCGGTCTTGATTGCACCTAACCGTCA3’ | attB2 reverse primer for cloning of MaLAR promoter into entry clone for Y1H assay |
| proMaANS_For | 5’GGGGACAAGTTTGTACAAAAAAGCAGGCTccCAGAATCAACAGTAAAAAAAAAGGAGG3’ | attB1 forward primer for cloning of MaANS promoter into entry clone for Y1H assay |
| proMaANS_Rev | 5’GGGGACCACTTTGTACAAGAAAGCTGGGTaTAGGAGAAGCTTCCGCTACTGGCAG3’ | attB2 reverse primer for cloning of MaANS promoter into entry clone for Y1H assay |
